# Beyond SOx: Mining Sulfur Bacteria Produce Less Acid due to Unidentified Enzyme Pathways

**DOI:** 10.64898/2025.12.01.691697

**Authors:** Jay Gordon

**Affiliations:** Civil and Mineral Engineering, University of Toronto, Toronto, Ontario, Canada

**Keywords:** SOx_1_, S_4_I_2_, *Halothiobacillus*_3_, *Thiomonas*_4_, *Pandoraea*_5_, sulfur_4_, mine wastewater_5_, acidity_6_

## Abstract

When thiosulfate is abundant, recent research indicates that the complete Sulfur Oxidation (cSOx) pathway is prevalent in oxic wastewater, while the incomplete Sulfur Oxidation and reverse Dissimilarity Sulfur Reduction (iSOx + rDSR) pathway has been found to be active under suboxic conditions, paired with nitrate reduction. However, both the activity of S_4_I (or other enzyme systems) and the suboxic sulfur reactions which occur when the rDSR pathway is not present in an SOB community remain unexplored. During this study, a series of 6 L benchtop microcosms were inoculated with *Halothiobacillus spp.*-dominated (containing cSOx and S_4_I pathways) sulfur oxidizing bacterial (SOB) communities preserved from an active tailing reservoir. The 16 microcosms were provided with high concentrations of the sulfur oxidation intermediate (SOI) compounds thiosulfate or tetrathionate under oxic and suboxic conditions to amplify the acid generating processes. The study identifies *Halothiobacillus*, *Thiomonas,* and *Pandoraea*, as the key SOB genera processing SOI. These microcosms demonstrate that protons yields resulting from thiosulfate oxidation by SOB are less than half of those predicted for thiosulfate oxidation, amongst the lowest reported for this genera. Despite adjustments for ion activity and PO_4_^2-^ buffering, this difference is indicative of mechanisms beyond cSOx. Further, when offered tetrathionate under oxic conditions, undescribed reactions unmatched to known sulfur enzymes were likely employed, since disproportionation is insufficient to explain observations. Meanwhile, suboxic depletion of thiosulfate, without changes in nitrate or nitrite concentrations sufficient to indicate their use as TEA requires further investigation.

## Introduction

Mine wastewater systems contain communities of sulfur-oxidizing bacteria (SOB) that influence the fate of sulfur oxidation intermediates (SOI) and resulting acidity (Whaley-Martin et al. 2019, 2023; Lopes et al. 2020; Miettinen et al. 2021; Gordon et al. 2024; Twible et al. 2024). Thiosalts (S_n_O_x_^2−^, common forms of SOI compounds containing combinations of sulfur and oxygen) are generated during milling processes and are a water quality concern for metal mines (Miranda-Trevino et al. 2013; Musuku et al. 2023). However, other than the field research mentioned above, few studies link a metagenomic understanding of sulfur oxidation with metal mine wastewater systems (SI Table 1). Instead, most genetic studies consider acidophilic genera and their contributions to acid rock drainage (ARD) or applications for bioleaching (Bugaytsova and Lindström 2004; Chen et al. 2004, 2015; Denef et al. 2010; Watling et al. 2014; Hua et al. 2015; Grettenberger et al. 2019; Wang et al. 2019; Camacho et al. 2020; Pakostova et al. 2022; Opara et al. 2023).

The SOB communities identified in circumneutral wastewater from metal mines processing sulfidic ores include key genera such as *Halothiobacillus, Thiomonas, Thiobacillus, Thiovirga, Sulfuricurvum, Sulfurovum,* and *Desulfurivibrio* (Whaley-Martin et al. 2019, 2023; Lopes et al. 2020; Miettinen et al. 2021; Gordon et al. 2024). These genera metabolically oxidize SOI via three sulfur oxidation pathways: the complete sulfur oxidation (cSOx), the incomplete SOx and reverse dissimilatory sulfur reduction (iSOx + rDSR), or the tetrathionate intermediate (S_4_I) (Friedrich et al. 2001; Ghosh and Dam 2009; Wasmund et al. 2017; Watanabe et al. 2019; Whaley-Martin et al. 2023a; Gordon et al. 2024a). The cSOx pathway is common in mine systems where thiosulfate is ubiquitous in processed tailings from sulfidic ore (equation 1):

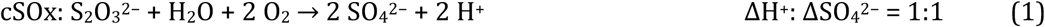

Geochemical studies focusing on sulfur oxidation in mine wastewater communities would suggest that the cSOx pathway is dominant in the oxic epilimnion of mine wastewater systems where the community also lacks the enzymes required to couple thiosulfate oxidation with nitrate reduction (Whaley-Martin et al. 2023a; Twible et al. 2024). Yet, as thiosulfate is consumed by *Halothiobacillus* and *Acidithiobacillus* in these systems, it fails to produce an observed ΔH^+^: ΔSO_4_^2−^ of 1:1 which would be expected from activity of the cSOx pathway (Bernier and Warren 2007; Warren et al. 2008; Whaley-Martin et al. 2019; Camacho et al. 2020b; Gordon et al. 2024). Whaley-Martin et al. (2019) isolated and cryopreserved communities from an active tailings impoundment which comprised ∼76–100 % *Halothiobacillus*. Because the flasks were open to the atmosphere, the SOB reactions were assumed to occur under oxic conditions, yet the ΔH^+^: ΔSO_4_^2−^ fell well below the theoretical cSOx ratio (Whaley-Martin et al. 2019). These SOB communities were used as inocula for this study to explore the phenomenon, perhaps due to S_4_I pathway activity, further.

While the detailed mechanisms of the cSOx pathway are well established, there is a lack of consensus about the enzymatic stages of the S_4_I pathway (SI Table 2). Only the first stage of the S_4_I pathway is universally agreed upon: the oxidative condensation of two thiosulfate molecules into tetrathionate facilitated by thiosulfate dehydrogenase (TsdA), or occasionally DoxDA, in wide range of genera such as *Paracoccus thiocyanatus*, *Allochromatium vinosum*, *Campylobacter jejuni*, *Erythrobacter flavus*, and *Acidithiobacillus ferrooxidans*, see equation 2 (Müller et al. 2004; Brito et al. 2015; Jenner et al. 2019; Rameez et al. 2020; Zhang et al. 2020; Yu et al. 2021; Du et al. 2022):

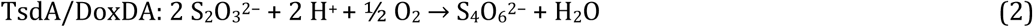

The widespread occurrence of *tsdA* across SOB genera, including *Halothiobacillus,* suggests that the S_4_I pathway often occurs in mine wastewater systems in both Canada and Spain, a mechanism that has been previously overlooked (Miettinen et al. 2021; Whaley-Martin et al. 2023). Following tetrathionate formation by TsdA, TetH and SoxB have both been proposed as mechanisms for the second stage, and it is an open question whether zero valent sulfur (ZVS) can form during the S_4_I pathway {SI Table 2 (Pyne et al. 2018; Cai et al. 2022)}.

This study used a series of 16 benchtop laboratory microcosms inoculated with SOB communities that had been cryogenically preserved from two depths of an active mine tailings impoundment (Whaley-Martin et al., 2019). The microcosms — grown with high density SOB populations, offered high concentrations of thiosulfate or tetrathionate, and monitored for changes in sulfur speciation and acidity generation — were designed to isolate and amplify the acid-generating phenomenon observed on site in field and mesocosm experiments (Whaley-Martin et al. 2023; Gordon et al. 2024). The experiments then explored (i) if acid and sulfate generation aligned with theoretical ratios for SOI oxidation, (ii) the roles of oxygen and nitrate as electron acceptors for sulfur oxidation, and (iii) if communities initially dominated by *Halothiobacillus* spp. have sulfur enzymes (a tetrathionate amendment targeted the S_4_I pathway) that can function alongside or instead of the cSOx pathway. By exploring aspects of microbial sulfur oxidation, this research seeks to understand how sulfur metabolism affects acidity generation in mine site wastewaters and to provide a theoretical basis for proactive management. To the best of our knowledge, only two other study explores a metagenomic understanding of sulfur metabolism of SOB communities isolated from circumneutral mine wastewaters, considering both their capacity for thiosalt oxidation and the geochemical products of their sulfur pathways (Twible et al. 2024; Liu et al. 2025).

## Methods

The following section describes the experimental design, sampling, analysis and calculations performed on mixed SOB cultures. The isolated SOB cultures were from the oxidation reservoir of an active tailings impoundment, cryopreserved, and regrown in benchtop microcosms.

### Benchtop 6 L Microcosms

A series of 16 benchtop microcosms were used to explore the S_4_I pathway reactions (under oxic and suboxic conditions) in *Halothiobacillus*- and *Thiomonas*-dominated communities using pH, sulfur speciation and 16S rRNA gene expression data. The microcosms were staged over three experiments (Experiments A, B, and C) for feasibility, and high concentrations of sulfur amendments were employed to exaggerate the acid-generating phenomenon observed on site. Experiments A, B, and C were performed on the laboratory benchtop in 6 L glass Erlenmeyer flasks (6000 mL Pyrex No. 1980), autoclaved for sterility, and filled to capacity with 6 L of SOB medium (0.1-μm filtered for sterility using ThermoScientific™ Nalgene™ Rapid-Flow™ Sterile Disposable Filter Units with PES membrane) containing 2.0 mM MgSO_4_, 0.2 mM NH_4_Cl, 1.4 mM K_2_HPO_4_, 8.3 mM NaNO_3_, and a trace element solution (Camacho, Jessen, et al. 2020) in all experiments (Fig 1). Microcosms were then inoculated with 2 or 10 m SOB communities, and fitted with HOBO data logger probes (sterilized with 70% vol/vol ethanol) for continuous DO and pH measurements (DO: HOBO® U26-001, and pH: HOBO® MX2501, Fig 1). Note, the pH loggers were used to log data every 60 seconds throughout all experiments. However, the dissolved oxygen (DO) data loggers were used to collect oxygen concentrations at t_0_ and t_end_ of Experiments A and B; in Experiment C, methodology was improved to allow DO-logging at 60-s intervals. Sample tubing (Crack-Rst Polyethylene-Lined EVA tubing for Food & Beverage, sterilized in 70% vol/vol ethanol) was then placed in the microcosms, and the tops were sealed with ethanol-sterilized Parafilm ® M P 7793.

**Fig 1.**
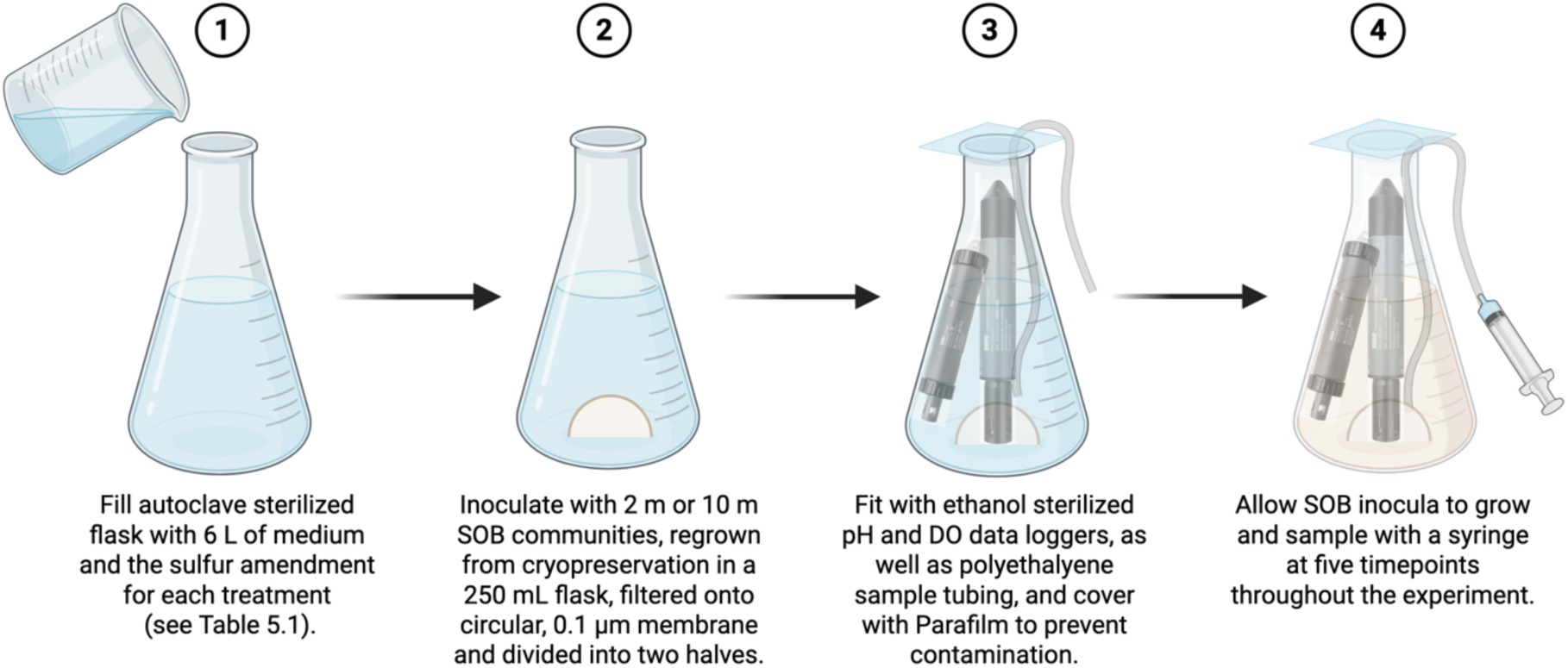
Microcosm setup for Experiments A, B, and C. The diagram above indicates the steps taken to assemble the probes, sample tubes and inocula in each of the 16 experimental microcosms. The HOBO DO data loggers were used to collect oxygen concentrations at t_0_ and t_end_ of Experiments A and B; in Experiment C, methodology was improved to allow DO-logging at 60-s intervals. (Figure created with BioRender.com.)

Sulfur amendments, which varied by experiment, were dissolved in Milli-Q ultrapure filtered water. Sulfur amendments were 8.0 mM S-S_4_O_6_^2−^ (as K_2_S_4_O_6_) in Experiments A and B and 11.4 mM S-S_2_O_3_^2−^ (as Na_2_S_2_O_3_) in Experiment C (Table 1). Microcosms were then inoculated with *Halothiobacillus*-dominated SOB communities collected from two depths of an oxidation reservoir on an active mine site in September 2017 (2 m and 10 m), cryopreserved, and regrown in similar media, resulting in treatments described in (Table 1).

**Table 1.** Experimental Treatments for 16 Benchtop microcosms.

| Experiment | Treatment | [SOI]<br>(mM) | [NO<sub>3</sub>]<br>(mM) | Rep |
| --- | --- | --- | --- | --- |
| A | Abiotic Control (no SOI) | - | 8.3 | - |
| A | Abiotic Control + Tetrathionate | 8.0 S-S<sub>4</sub>O<sub>6</sub><sup>2-</sup> | 8.3 | - |
| A | 2 m SOB (No SOI) | - | 8.3 | R1 |
| A | 2 m SOB (No SOI) | - | 8.3 | R2 |
| A | 2 m SOB + Tetrathionate | 8.0 S-S<sub>4</sub>O<sub>6</sub><sup>2-</sup> | 8.3 | R1 |
| A | 2 m SOB + Tetrathionate | 8.0 S-S<sub>4</sub>O<sub>6</sub><sup>2-</sup> | 8.3 | R2 |
| B | 10 m SOB (No SOI) | - | 8.3 | R1 |
| B | 10 m SOB (No SOI) | - | 8.3 | R2 |
| B | 10 m SOB + Tetrathionate | 8.0 S-S<sub>4</sub>O<sub>6</sub><sup>2-</sup> | 8.3 | R1 |
| B | 10 m SOB + Tetrathionate | 8.0 S-S<sub>4</sub>O<sub>6</sub><sup>2-</sup> | 8.3 | R2 |
| C | 2 m SOB + Thiosulfate | 11.4 S-S<sub>2</sub>O<sub>3</sub><sup>2-</sup> | 8.3 | R1 |
| C | 2 m SOB + Thiosulfate | 11.4 S-S<sub>2</sub>O<sub>3</sub><sup>2-</sup> | 8.3 | R2 |
| C | 10 m SOB + Thiosulfate | 11.4 S-S<sub>2</sub>O<sub>3</sub><sup>2-</sup> | 8.3 | R1 |
| C | 10 m SOB + Thiosulfate | 11.4 S-S<sub>2</sub>O<sub>3</sub><sup>2-</sup> | 8.3 | R2 |
| C | Abiotic Control + Thiosulfate | 11.4 S-S<sub>2</sub>O<sub>3</sub><sup>2-</sup> | 8.3 | R1 |
| C | Abiotic Control + Thiosulfate | 11.4 S-S<sub>2</sub>O<sub>3</sub><sup>2-</sup> | 8.3 | R2 |

(-) indicates no amendment of this type added to the mesocosm.

These *Halothiobacillus*-dominated communities were similar to those used to inoculate the 500 L mesocosms, but were collected from the tailings impoundment and cryopreserved the previous year. A few days prior to the initiation of each experiment, the cryopreserved SOB community was resuscitated by thawing and placing it into a 500 mL, autoclave-sterilized glass flask filled with 500 mL of medium (recipe above, amended with either thiosulfate or tetrathionate to match the experimental treatment). To inoculate the microcosms with these communities, cultures were filtered onto 0.1-μm paper filters using sterile filter columns (ThermoScientific™ Nalgene™, see above). The filter paper with microbial biofilm was cut in two, and one semi-circle was transferred into each of the two replicates of the sterile microcosms using tweezers (sterilized in 70% vol/vol ethanol for two minutes) inside a biological safety cabinet.

In Experiment C, where suboxic conditions were encountered, two additional tests for DO diffusion into the microcosms were performed. First, a replicate of the 6 L flask was filled with 5 L of thiosulfate-amended medium and inoculated with an SOB community. Once suboxic conditions were reached at the normal probe depth (10 cm from the base of the 6 L flask), DO concentrations were measured at 20 cm from the base and <1 cm below the surface of the media to ensure that probe readings at these depths were also suboxic. Second, a sterile flask filled with MilliQ water was sparged with N_2_ (g) for 25 min to achieve suboxia, and a DO data logger was used to check for oxygen ingress from atmospheric diffusion over 24 h (compared against a control of open water).

### Sample Collection and Geochemical Analysis

Samples for sulfur speciation (S^0^, S_2_O_3_^2−^, SO_3_^2−^, SO_4_^2−^, and Total S) were collected for five timepoints (t_0_, t_1_, t_2_, t_3_, t_4_, and t_end_) in all Experiments (A, B, and C); samples for HS^−^ concentrations were collected for all timepoints in Experiments A and B, but could not be measured in Experiment C because of the high biomass. Sampling frequency was decreased in Experiment C as acid generation occurred less rapidly. Samples for nitrogen speciation (NO_2_^−^, NO_3_^−^, and NH_4_^+^) were taken at the timepoints in Experiment C where suboxic conditions were established.

In all benchtop experiments, samples were collected using a syringe (Fisherbrand 60 mL plastic syringe, rinsed with water from the mesocosm prior to sampling) to draw water from the microcosms via silicon tubing (Crack-Rst Polyethylene-Lined EVA tubing for Food & Beverage, ¼″). Samples were filtered, when necessary, with a 0.2-μm tip and immediately preserved in the fridge or freezer; ion chromatograph samples were 0.2-μm filtered; ICP-OES samples were unfiltered and acid preserved (0.02% HNO_3_); HPLC samples were unfiltered and frozen. Samples for thiosulfate and sulfite measurements by HPLC were frozen after monobromobimane derivatization of unfiltered samples (see sections 2.3 and 2.4.1). Concentrations of sulfur and nitrogen species (sulfate, nitrate, nitrite, and ammonia), were measured in 0.2-μm filtered samples using Thermo Scientific Dionex ICS-6000 for ion chromatography (see sections 2.3 and 2.4.1). Shimadzu LC-20AD Prominence high performance liquid chromatography was used to measure thiosulfate and sulfite {monobromobimane derivatization method (Rethmeier et al. 1997)} and elemental sulfur (chloroform extraction method); and a ThermoFisher Scientific inductively coupled plasma–optical emission spectrograph (ICP-OES) was employed for total sulfur analyses on, using detection wavelength of 180.731 nm radial. Samples for S^0^ analysis for Experiments A and B were performed according to the methods outlined in Whaley Martin et al (2020) and Yan et al (2022). However, after processing the first replicate of S^0^ samples from Experiment C, HPLC performance was improved by adding a 20-min sonication step, following redissolving the extracted S^0^ in chloroform, prior to filtering through 0.2-µm PTFE tips. This resulted in a method that was only semi-quantitative. Sulfide samples were quantified using a HACH spectrophotometer using the Methylene Blue Method 8131.

### DNA Extraction, Quantification, Sequencing and Analyses

DNA was collected by filtering 0.5 L of water from the microcosms through 0.1-μm filters using sterile filter columns (Thermo Scientific™ Nalgene™ Rapid-Flow™ Sterile Disposable Filter Units with CN Membrane) and immediately excising and freezing the filter paper. Water was transferred into filter columns using a syphon, and DNA extractions were performed in a UV-sterilized biological safety cabinet using the Qiagen’s DNeasy PowerWater DNA Isolation Kit using the standard protocol.

Illumina MiSeq region V4 amplicon sequencing was performed at McMaster University’s Metagenomics Facility (Hamilton, Ontario) as described in section 3.2.8 above. Once extracted, the DNA were quantified and the 515F/806R primer set was used to amplify the 16S V4 region and dual-indexed Illumina adapters were added to the amplicons. The amplicons were then quality-assessed via agarose gel electrophoresis before being normalized using a SequelPrep normalization kit. The resulting sequences were filtered and trimmed using Cutadapt (with a minimum “Q30” Phred quality score, indicating that likelihood of an incorrect base call is less than 1/1000). Fastq read files were filtered to a trimmed length (240 bp in the forward direction and 160 bp in the reverse direction) and processed according to the DADA2 pipeline (version 3.1.6) to remove bimeras and chloroplast or mitochondrial sequences. Taxonomy was then assigned using to SILVA database (version 138.1) and R (version 4.4.4) in RStudio (version 1.2).

### Proton Yield Calculations

Hydrogen ion concentrations ([H^+^]) and their changes with time [ΔH^+^], were calculated according to equations 3:

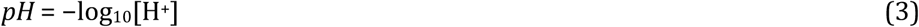

This simplification is accurate within the range of 80 - 100 %, as:

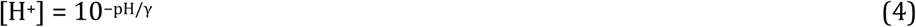

The ion activity coefficient (γ±) was estimated for the solution with a low ionic strength (I <0.1 M) of I ∼ 0.03-0.05 molal (given a media of 2.0 mM MgSO_4_, 0.2 mM NH_4_Cl, 1.4 mM K_2_HPO_4_, 8.3 mM NaNO_3_, and 2.0 mM S_4_O_6_^2-^ or 5.7 mM S_2_O_3_^2−^), according to the Davis equation, equation 5:

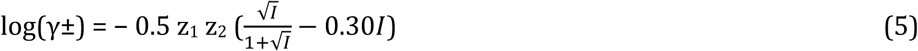

Where z_1_ and z_2_ are ion charges, and I is ionic strength, and 0.5 is the temperature dependant coefficient (A) at 25°C (Stumm and Morgan 1995). This produces an activity coefficient of ∼0.82-0.85. The range of 80 - 100 % is represented as field of potential concentrations in figure 3.

As a result, the conversion allows for a stoichiometric exploration of H^+^ - S ratios, accurate within the range of 80 - 100 %, as per equation 6.

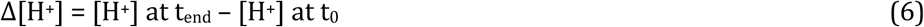

To account for the magnitude of phosphate and carbonate buffering in the biotic systems, the buffering capacity was calculated. Based on an initial concentration of 1.4 mM K_2_HPO_4_ in the media at pH ∼7, and the [HPO_4_^2−^] / [H_2_PO_4_^−^] = 1.38 ratio (from the pKa = 6.86), the concentrations of the acid (H_2_PO_4_^−^) would theoretically be 0.6 mM, and the conjugate base (HPO_4_^2−^) would be 0.8 mM at the beginning of the microcosm experiments. Since 1 mole of HPO_4_^2−^ can neutralize 1 mole H^+^, this would mean that 0.8 mM H^+^ could be neutralized in the media before the buffer is fully exhausted. When incorporating similar calculations for carbonate buffering, the total buffering capacity (β) of the media ranged from β = 0.01 - 2.8 (VMinteq version 4.1 calcs, Dr. Simon Apte, personal correspondence):

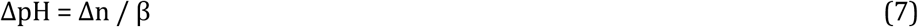

Where Δn is the number of moles of an acid added to solution; β is buffer capacity; ΔpH is the difference between the initial pH of the buffer and the pH of the buffer after the addition.

The VMinteq modeling was performed in version 4.1 by assuming a gas equilibrium with atmospheric CO_2_ (a function under the menu “gases”) and calculating buffering via a mass balance at 22°C (in correspondence with Simon Apte). The resulting calculations for buffering in the media indicated that this buffering would produce gradual pH changes above pH 6.8 (when [HPO_4_^2−^] / [H_2_PO_4_^−^] =1) would be expected, accelerating slightly to 5.5 (Fig D1). After this point the buffer should be fully exhausted, and the pH can be expected to drop sharply if acid generation continues (Fig D1).

Therefore, the Δ[H^+^] value used in figure 4 was calculated using VMinteq outputs as a single value. This VMinteq output would be equal to approximately (equation 8):

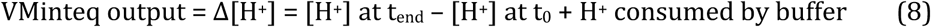

The Δ[H^+^] was then used to calculate a proton yield by mole of sulfate produced through SOI oxidation ( ΔH^+^ : ΔSO_4_^2−^ ).

### Statistical and Data Analyses

Statistical analyses were carried out in R version 4.4.4, utilizing the Vegan package version 2.6.4. The analyses were carried out on triplicate samples for geochemistry, unless noted, with standard deviation calculated on mean analytical data from triplicate injections. “Less than detection limit” chemical or biological data were treated as zero for statistical analyses. Data were displayed using Ggplot, and the methods figure was created in BioRender.

Treatments were frequently performed in duplicate. Although low replicate sizes limit the strength of statistical analysis, a duplication of trends in sulfur speciation or pH changes provided some reassurance that observed phenomena were a result or treatments and not random variability. Where replicates showed a unique behaviour (example: Fig 3, Experiment C), this was interpreted as due to differences in the dominate microbial community.

## Results

A summary of the geochemical data can be found in Table 2, and the complete dataset (sulfur speciation, nitrogen speciation, oxygen and pH) in SI Table 3. Table 2 summarizes t_0_ and t_end_ data from the 16 microcosms, with a sulfur mass balance calculated from the sulfur species quantified: [HS^−^], [S^0^], [S-S_2_O_3_^2−^], [SO_3_^2−^], and [SO_4_^2−^]. The standard error from the tetrathionate (S_4_O_6_^2−^) analysis was too high for reliable reporting; therefore, these data were omitted from the geochemical dataset. Although this limits the exploration of the later stages of the S_4_I pathway, changes in the remaining sulfur species nonetheless produced interesting findings, described below.

**Table 2.**
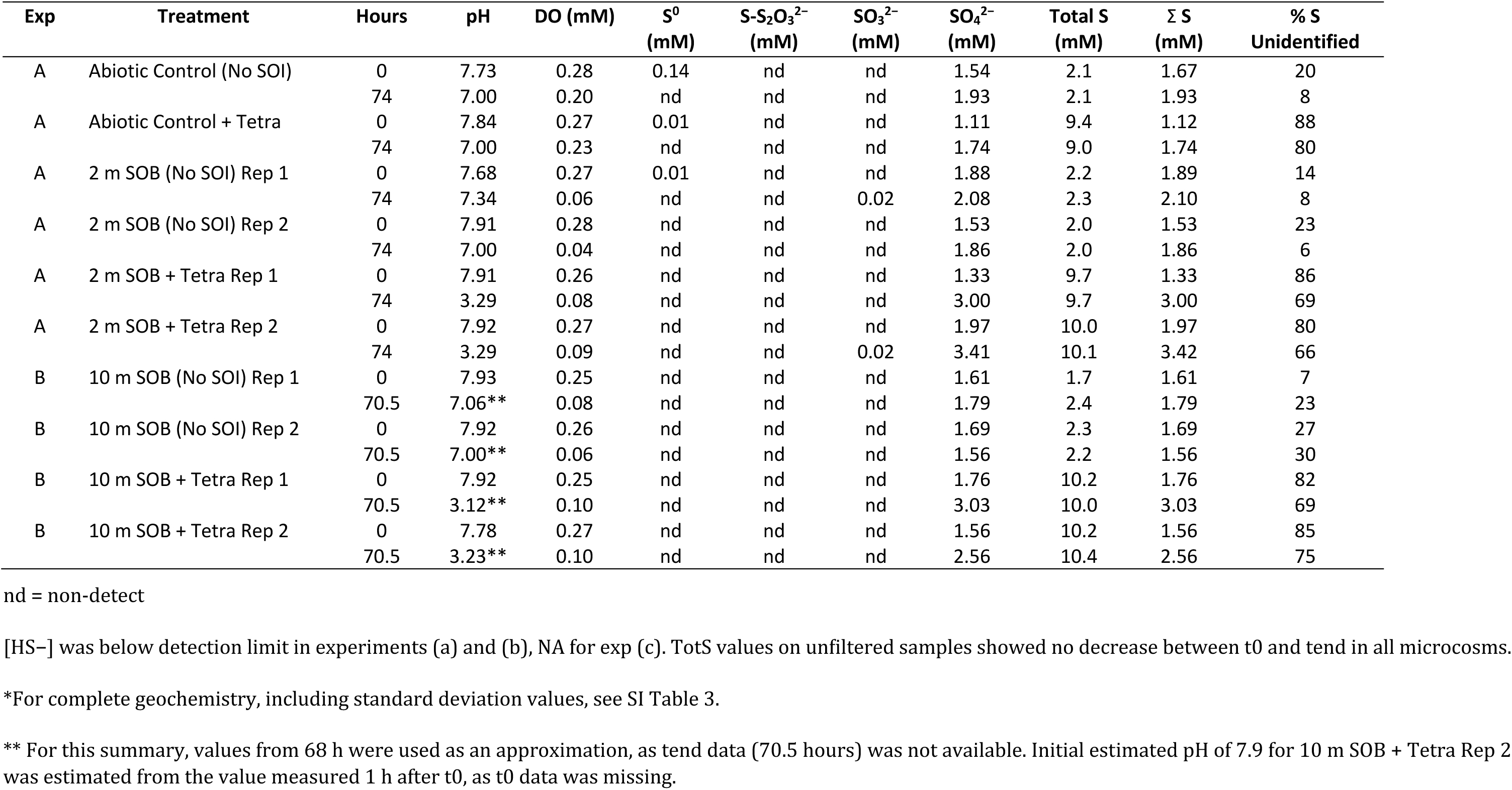
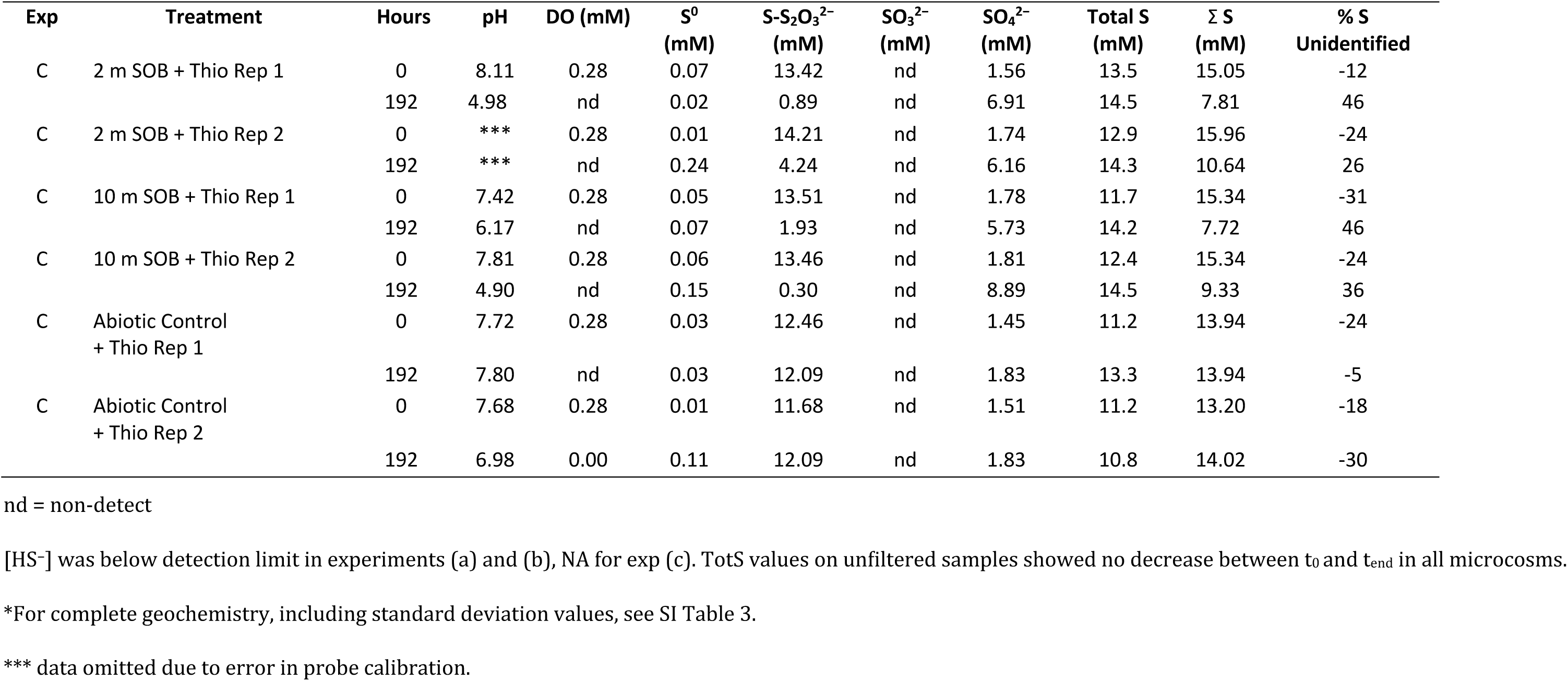
Summary of Geochemical Data from 16 Microcosms*.

| Exp | Treatment | Hours | pH | DO (mM) | S<sup>0</sup><br>(mM) | S-S<sub>2</sub>O<sub>3</sub><sup>2-</sup><br>(mM) | SO<sub>3</sub><sup>2-</sup><br>(mM) | SO<sub>4</sub><sup>2-</sup><br>(mM) | Total S<br>(mM) | Σ S<br>(mM) | % S<br>Unidentified |
| --- | --- | --- | --- | --- | --- | --- | --- | --- | --- | --- | --- |
| A | Abiotic Control (No SOI) | 0 | 7.73 | 0.28 | 0.14 | nd | nd | 1.54 | 2.1 | 1.67 | 20 |
|  |  | 74 | 7.00 | 0.20 | nd | nd | nd | 1.93 | 2.1 | 1.93 | 8 |
| A | Abiotic Control + Tetra | 0 | 7.84 | 0.27 | 0.01 | nd | nd | 1.11 | 9.4 | 1.12 | 88 |
|  |  | 74 | 7.00 | 0.23 | nd | nd | nd | 1.74 | 9.0 | 1.74 | 80 |
| A | 2 m SOB (No SOI) Rep 1 | 0 | 7.68 | 0.27 | 0.01 | nd | nd | 1.88 | 2.2 | 1.89 | 14 |
|  |  | 74 | 7.34 | 0.06 | nd | nd | 0.02 | 2.08 | 2.3 | 2.10 | 8 |
| A | 2 m SOB (No SOI) Rep 2 | 0 | 7.91 | 0.28 | nd | nd | nd | 1.53 | 2.0 | 1.53 | 23 |
|  |  | 74 | 7.00 | 0.04 | nd | nd | nd | 1.86 | 2.0 | 1.86 | 6 |
| A | 2 m SOB + Tetra Rep 1 | 0 | 7.91 | 0.26 | nd | nd | nd | 1.33 | 9.7 | 1.33 | 86 |
|  |  | 74 | 3.29 | 0.08 | nd | nd | nd | 3.00 | 9.7 | 3.00 | 69 |
| A | 2 m SOB + Tetra Rep 2 | 0 | 7.92 | 0.27 | nd | nd | nd | 1.97 | 10.0 | 1.97 | 80 |
|  |  | 74 | 3.29 | 0.09 | nd | nd | 0.02 | 3.41 | 10.1 | 3.42 | 66 |
| B | 10 m SOB (No SOI) Rep 1 | 0 | 7.93 | 0.25 | nd | nd | nd | 1.61 | 1.7 | 1.61 | 7 |
|  |  | 70.5 | 7.06** | 0.08 | nd | nd | nd | 1.79 | 2.4 | 1.79 | 23 |
| B | 10 m SOB (No SOI) Rep 2 | 0 | 7.92 | 0.26 | nd | nd | nd | 1.69 | 2.3 | 1.69 | 27 |
|  |  | 70.5 | 7.00** | 0.06 | nd | nd | nd | 1.56 | 2.2 | 1.56 | 30 |
| B | 10 m SOB + Tetra Rep 1 | 0 | 7.92 | 0.25 | nd | nd | nd | 1.76 | 10.2 | 1.76 | 82 |
|  |  | 70.5 | 3.12** | 0.10 | nd | nd | nd | 3.03 | 10.0 | 3.03 | 69 |
| B | 10 m SOB + Tetra Rep 2 | 0 | 7.78 | 0.27 | nd | nd | nd | 1.56 | 10.2 | 1.56 | 85 |
|  |  | 70.5 | 3.23** | 0.10 | nd | nd | nd | 2.56 | 10.4 | 2.56 | 75 |

**Table 2 cont.** Geochemical Summary from 16 Microcosms\*
| Exp | Treatment | Hours | pH | DO (mM) | S<sup>0</sup><br>(mM) | S-S<sub>2</sub>O<sub>3</sub><sup>2-</sup><br>(mM) | SO<sub>3</sub><sup>2-</sup><br>(mM) | SO<sub>4</sub><sup>2-</sup><br>(mM) | Total S<br>(mM) | Σ S<br>(mM) | % S<br>Unidentified |
| --- | --- | --- | --- | --- | --- | --- | --- | --- | --- | --- | --- |
| C | 2 m SOB + Thio Rep 1 | 0 | 8.11 | 0.28 | 0.07 | 13.42 | nd | 1.56 | 13.5 | 15.05 | -12 |
|  |  | 192 | 4.98 | nd | 0.02 | 0.89 | nd | 6.91 | 14.5 | 7.81 | 46 |
| C | 2 m SOB + Thio Rep 2 | 0 | *** | 0.28 | 0.01 | 14.21 | nd | 1.74 | 12.9 | 15.96 | -24 |
|  |  | 192 | *** | nd | 0.24 | 4.24 | nd | 6.16 | 14.3 | 10.64 | 26 |
| C | 10 m SOB + Thio Rep 1 | 0 | 7.42 | 0.28 | 0.05 | 13.51 | nd | 1.78 | 11.7 | 15.34 | -31 |
|  |  | 192 | 6.17 | nd | 0.07 | 1.93 | nd | 5.73 | 14.2 | 7.72 | 46 |
| C | 10 m SOB + Thio Rep 2 | 0 | 7.81 | 0.28 | 0.06 | 13.46 | nd | 1.81 | 12.4 | 15.34 | -24 |
|  |  | 192 | 4.90 | nd | 0.15 | 0.30 | nd | 8.89 | 14.5 | 9.33 | 36 |
| C | Abiotic Control<br>+ Thio Rep 1 | 0 | 7.72 | 0.28 | 0.03 | 12.46 | nd | 1.45 | 11.2 | 13.94 | -24 |
|  |  | 192 | 7.80 | nd | 0.03 | 12.09 | nd | 1.83 | 13.3 | 13.94 | -5 |
| C | Abiotic Control<br>+ Thio Rep 2 | 0 | 7.68 | 0.28 | 0.01 | 11.68 | nd | 1.51 | 11.2 | 13.20 | -18 |
|  |  | 192 | 6.98 | 0.00 | 0.11 | 12.09 | nd | 1.83 | 10.8 | 14.02 | -30 |

nd = non-detect
[HS<sup>-</sup>] was below detection limit in experiments (a) and (b), NA for exp (c). TotS values on unfiltered samples showed no decrease between t<sub>0</sub> and t<sub>end</sub> in all microcosms.
\*For complete geochemistry, including standard deviation values, see SI Table 3.
\*\*\* data omitted due to error in probe calibration.

### Sulfur Speciation Patterns

In the four tetrathionate-amended microcosms with SOB (Experiments A and B), sulfate concentrations increased consistently (mean ΔSO_4_^2−^_2m_ = 1.55 ± 0.29 mM; mean ΔSO_4_^2−^_10m_ = 1.14 ± 0.55 mM, Fig 1, SI Table 3). There was some variability between reps in the speed of sulfate formation, with an early spike in sulfate concentration detected in one of the 10 m SOB communities. Yet, the overall increases of [SO_4_^2−^] were in the same order of magnitude across replicates, accounting for <20% of the 8.0 mM ΔS-S_4_O_6_^2−^ amendment (Fig 1, SI Table 3). This suggests either tetrathionate (S_4_O_6_^2−^) remained unmetabolized, or other polythionates (e.g., S_3_O_6_^2−^, S_5_O_6_^2−^) formed during tetrathionate oxidation. In addition to sulfate formation, small increases in thiosulfate were measured at the midpoints of the tetrathionate-amended microcosms from both depths (mean ΔS-S_2_O_3_^2−^_2m_ = 0.24 ± 0.01 mM; mean ΔS-S_2_O_3_^2−^_10m_ = 0.28 ± 0.01 mM, Fig 1). Yet, at its highest concentrations, molar S-S_2_O_3_^2−^ evolution accounted for <0.04% of the 8.0 mM S-S_4_O_6_^2−^ amendments, not substantially change the fraction of detected sulfur species. Tetrathionate data was of too low quality to be included in the analysis.

A sulfur mass balance {[TotS] – Σ([HS^−^], [S^0^], [S-S_2_O_3_^2−^], [SO_3_^2−^], [SO_4_^2−^]); mM} demonstrated that >50% of the S remained unaccounted for in both biotic and abiotic tetrathionate-amended microcosms (Experiments A and B; Table 2, SI Table 3). No substantial [HS^−^], [S^0^], or [SO_3_^2−^] were detected. This gap in the S mass balance suggests that other reactive sulfur compounds (S_react_) (Whaley-Martin et al. 2020) were present, likely as tetrathionate (S_4_O_6_^2−^) and possibly as other polythionates (e.g., S_3_O_6_^2−^, S_5_O_6_^2−^).

In the thiosulfate-amended microcosms with SOB in Experiment C, sulfate increases occurred at a consistent rate of ∼0.03 mM/h under both oxic (mean ΔSO_4_^2−^_2m_ = 0.8 ± 0.4 mM and mean ΔSO_4_^2−^_10m_ = 0.6 ± 0.3 mM between hours 0–24) and suboxic (mean ΔSO_4_^2−^_2m_ = 3.8 ± 0.5 mM; mean ΔSO_4_^2−^_10m_ = 4.3 ± 0.7 mM between hours 74–192) conditions (Fig 1, SI Table 3). In sharp contrast to the tetrathionate-amended microcosms, the thiosulfate concentrations in the thiosulfate-amended microcosms decreased by 11.2 ± 1.2 mM S-S_2_O_3_^2−^ (2 m SOB) and 12.4 ± 0.2 mM S-S_2_O_3_^2−^ (10 m SOB) over the course of the experiment, accelerating from 0.02 mM/h under oxic conditions to a mean rate of 0.09 mM/h after the systems became suboxic (Fig 1). While the microcosms were under oxic conditions (between 0-24 hours), thiosulfate depletion may have been equal to the rate of sulfate formation (mean -ΔS_2_O_3_^2−^_2m_ = 0.6 ± 0.9 mM and mean -ΔS_2_O_3_^2−^_10m_ = 0.4 ± 0.4 mM, SI Table 3). However, once suboxic conditions were reached, only 30–40% of the thiosulfate loss could be accounted for by sulfate formation, and the difference was not accounted for by ZVS formation.

For the thiosulfate-amended treatments, the sulfur mass balance {[TotS] = Σ ([S^0^], [S-S_2_O_3_^2−^], [SO_3_^2−^], [SO_4_^2−^]); as [HS^−^] was not quantifiable due to high biomass} indicates that all S could initially be accounted for (112–130% of TotS accounted for at t_0_ on unfiltered sample) as the measured sulfur species. During both the oxic and anoxic portions of Experiment C, low [S^0^] was detected in all microcosms (semi-quantitative measurements, SI Table 3). There was no SO_3_^2−^ detected in any of the systems. However, as the experiments progressed, a gap in the accounted-for S developed (24–75% S was not accounted for at t_end_), likely due to the formation of polythionates. A comparison of [TotS] at t_0_ and t_end_ indicates that no sulfur left the solution as H_2_S (g).

No statistically significant changes in sulfate concentration occurred in the abiotic controls amended with thiosulfate in Experiment C, nor in the biotic controls lacking SOI amendments (Table 2, SI Table 3). However, small increases in [SO_4_^2−^] were detected in the tetrathionate-amended abiotic controls in experiment a (ΔSO_4_^2−^_M2_ = 0.6 ± 0.4 mM, Table 2). These increases in [SO_4_^2−^] may be attributed to tetrathionate instability above pH 7, as the abiotic tetrathionate control remained between pH 7.7 and 7 over the course of the experiment; above pH 7, tetrathionate degrades abiotically into tri- and pentathionate, and eventually to thiosulfate and sulfate (Varga and Horváth 2007).

### pH Changes and Proton Yield from Sulfur Oxidation

Considering the pH data alone, different responses clearly arise from the combinations of SOB and sulfur substrate present each treatment. In the first two tetrathionate-amended experiments (A and B), acidity generation clearly resulted from tetrathionate metabolism by the SOB community (pH decreased from 7.9 to 3.1–3.3, Fig 2). The trajectory of acid generation was similar between 2 and 10 m communities, although slightly more rapid in the microcosms inoculated with SOB from the greater depth (experiment B, Fig 2). In contrast, in both biotic and abiotic controls exhibited only slight pH decreases (<1 pH unit; Fig 2), which can be accounted for by the acid-forming processes of the carbonate buffering system (Stumm and Morgan 1995), a slightly depressed by the phosphate buffering present (see methods).

**Fig 2.**
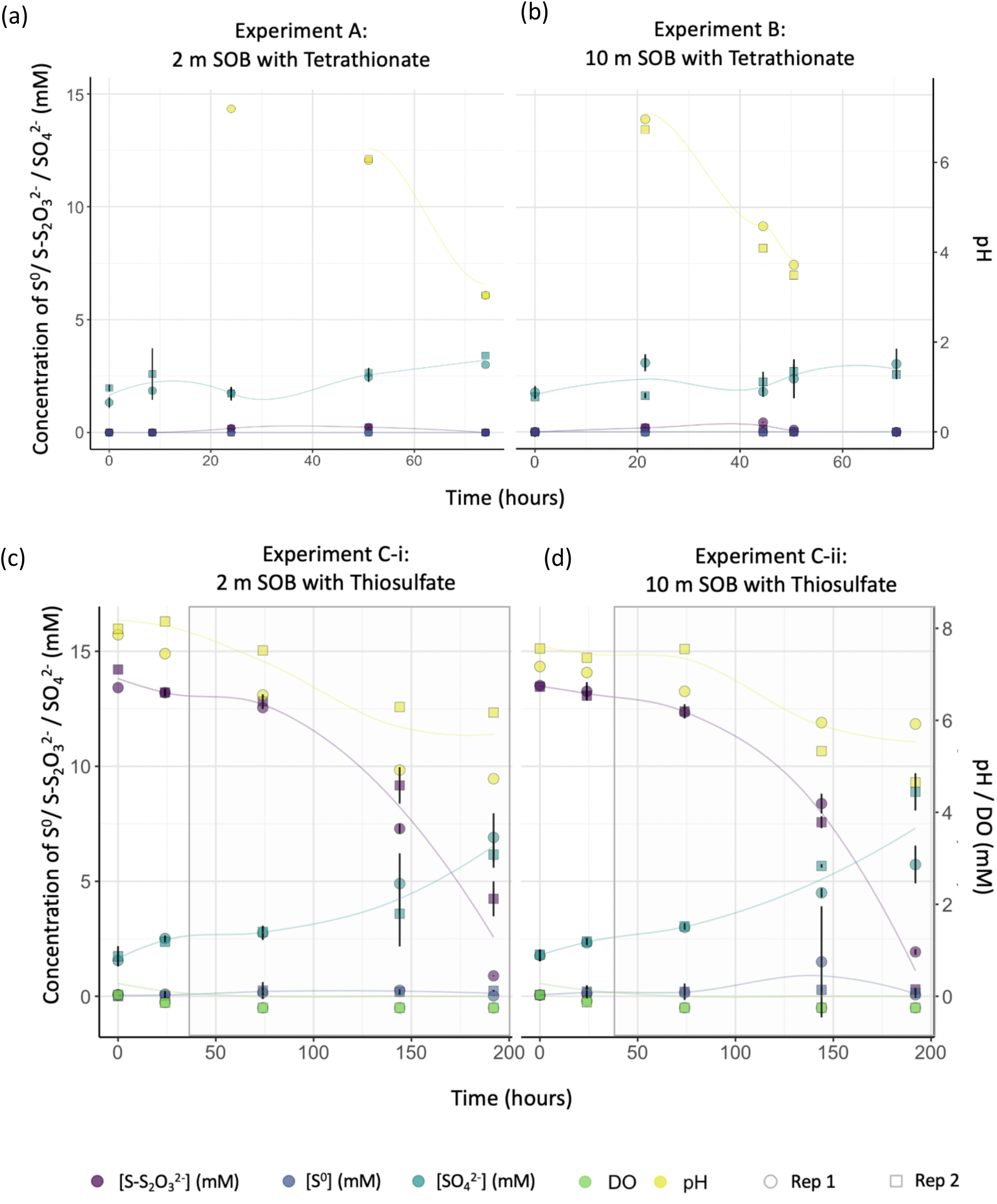
Changes in sulfur speciation (S^0^, S-S_2_O_3_^2−^, SO_4_^2−^) occurred in the 8 microcosms treated with sulfur-oxidation intermediate amendments and sulfur-oxidizing bacterial (SOB) communities. Samples were collected from two replicates (circles and squares and blue; some circles not visible due to overlap) of each of the following treatments: (a) 2 m SOB treated with 8.0 mM S-S_4_O_6_^2−^, (b) 10 m SOB treated with 8.0 mM S-S_4_O_6_^2−^, (c) 2 m SOB treated with 11.4 mM S-S_2_O_3_^2−^, and (c) 10 m SOB treated with 11.4 mM S-S_2_O_3_^2−^. Grey rectangles indicate suboxic portion of Experiment C. The [SO_3_^2−^] < limit of detection (LoD) for all experiments, and [HS−] < LoD for Experiments A and B.

**Fig 3.**
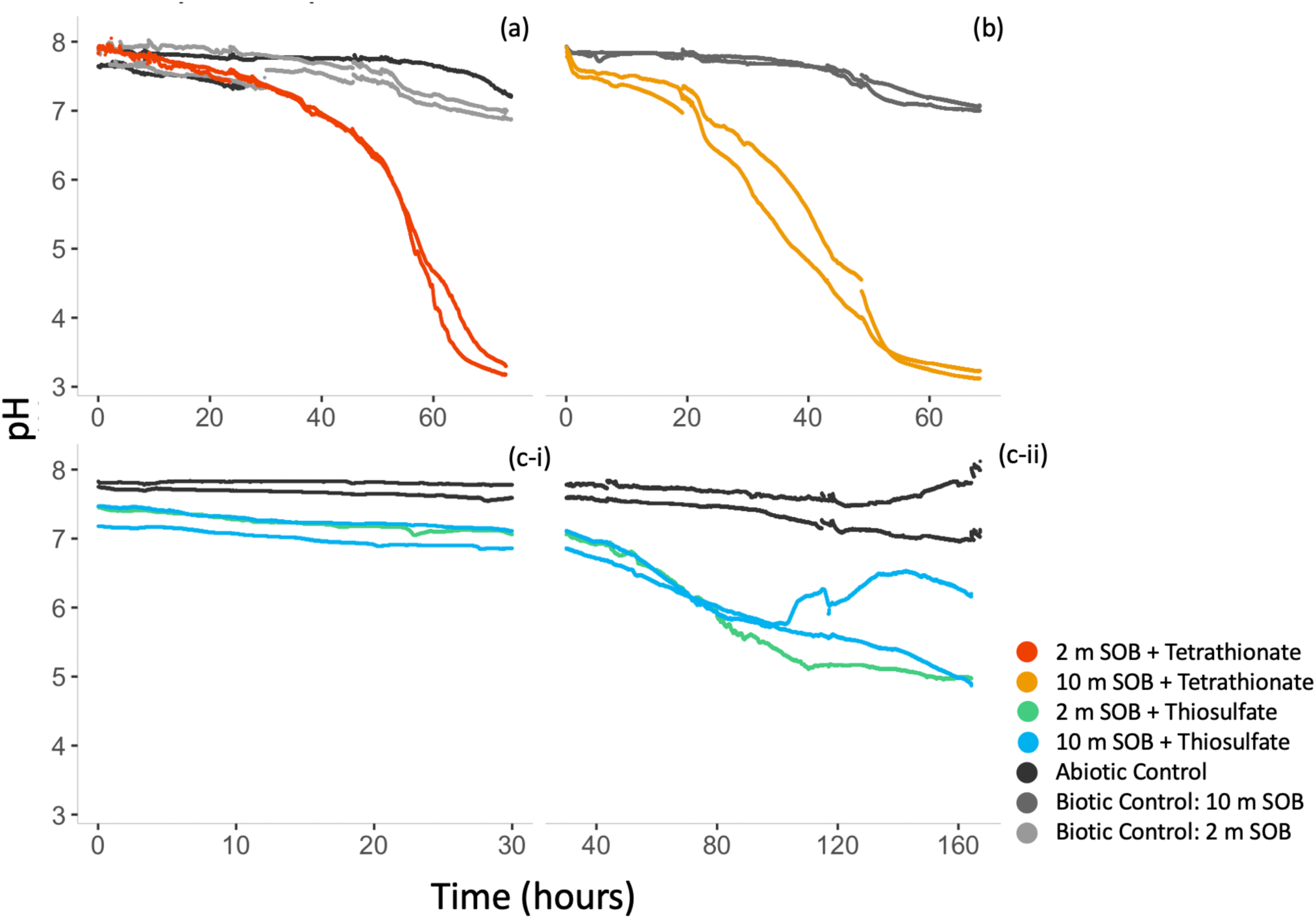
Acid generation by sulfur-oxidizing communities in (a) Ox Res 2 m SOB community with tetrathionate amendments (oxic) over 70 h, (b) Ox Res 10 m SOB community with tetrathionate amendments (oxic) over 70 h, and (c-i) oxic or (c-ii) suboxic portions of the experiment with 2 m and 10 m SOB communities from a tailings reservoir with thiosulfate amendments over 180 h. (Note: pH readings from 2 m SOB + Thiosulfate Replicate 2 were omitted because of a calibration error; however, a pH decrease was also observed.)

In the thiosulfate-amended Experiment C, acidity generation also clearly resulted from thiosulfate metabolism. The pH decreased from 7.4–8.1 to 5.0–6.2 in the microcosms which received both thiosulfate and SOB amendments, with most acid generated under suboxic conditions (Fig 2). Again, all controls only exhibited slight pH decreases due to carbonate buffering. However, in suboxic portion of this experiment, variation was observed in one of the 10 m SOB replicates. Unlike the other two microcosms, the pH in this community experienced a point of inflection at 100 h, when the system (at a pH of 5.6) switched from acidity-generating to -consuming, suggesting a shift in thiosulfate oxidation processes (Fig 2c).

Along with acid generation ([H^+^] increases), sulfate concentrations ([SO_4_^2−^]) also increased in the in the SOI-amended microcosms inoculated with SOB. In the tetrathionate-amended microcosms, the ΔH^+^ : ΔSO_4_^2−^ between t_0_ and t_end_ (0.33:1 for 2 m SOB, and 0.17:1 for 10 m SOB, see methods section for details of the calculation) was an order of magnitude below a theoretical 3:2 ratio for direct tetrathionate oxidation with oxygen (equation 3, Fig 3):

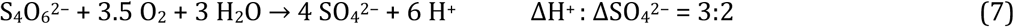

Even given the potential 20 % underestimation of [H^+^] resulting from the ion activity coefficient when converting pH data to directly proton concentration (see methods), the detected [H^+^] calculated from pH are low. In the tetrathionate amended systems, this difference can be accounted for by the carbonate and phosphate buffering in the system; when accounted for the in the [H^+^] calculations (see methods), the ΔH^+^ : ΔSO^42−^ is found to be within (for 2 m SOB), or slightly above (for 10 m SOB) the expected range for direct tetrathionate oxidation to sulfate (Fig 3).

However, in the thiosulfate-amended microcosms, the discrepancy between theoretical and actual was pronounced. Here, the ΔH^+^ : ΔSO_4_^2−^ between t_0_ and t_end_ was less than half the 1:1 theoretically predicted by direct thiosulfate oxidation to sulfate (equation 1, see Fig 3), even accounting for proton neutralization due to buffering. At less than half of the ΔH^+^ that would be expected via complete thiosulfate oxidation to sulfate, this observation cannot be explained by the stoichiometry of the cSOx pathway.

### Microbial Community Signals

At the beginning of each experiment (t_0_), biotic treatments were inoculated with communities isolated from either 2 m or 10 m depth from the tailings impoundment. Three key SOB genera were identified in these communities by referencing metagenomes previously sequenced from the oxidation reservoir and the current literature: *Halothiobacillus, Pandoraea,* and *Thiomonas* (Fig 4, SI Tables 4 and 5).

**Fig 4.**
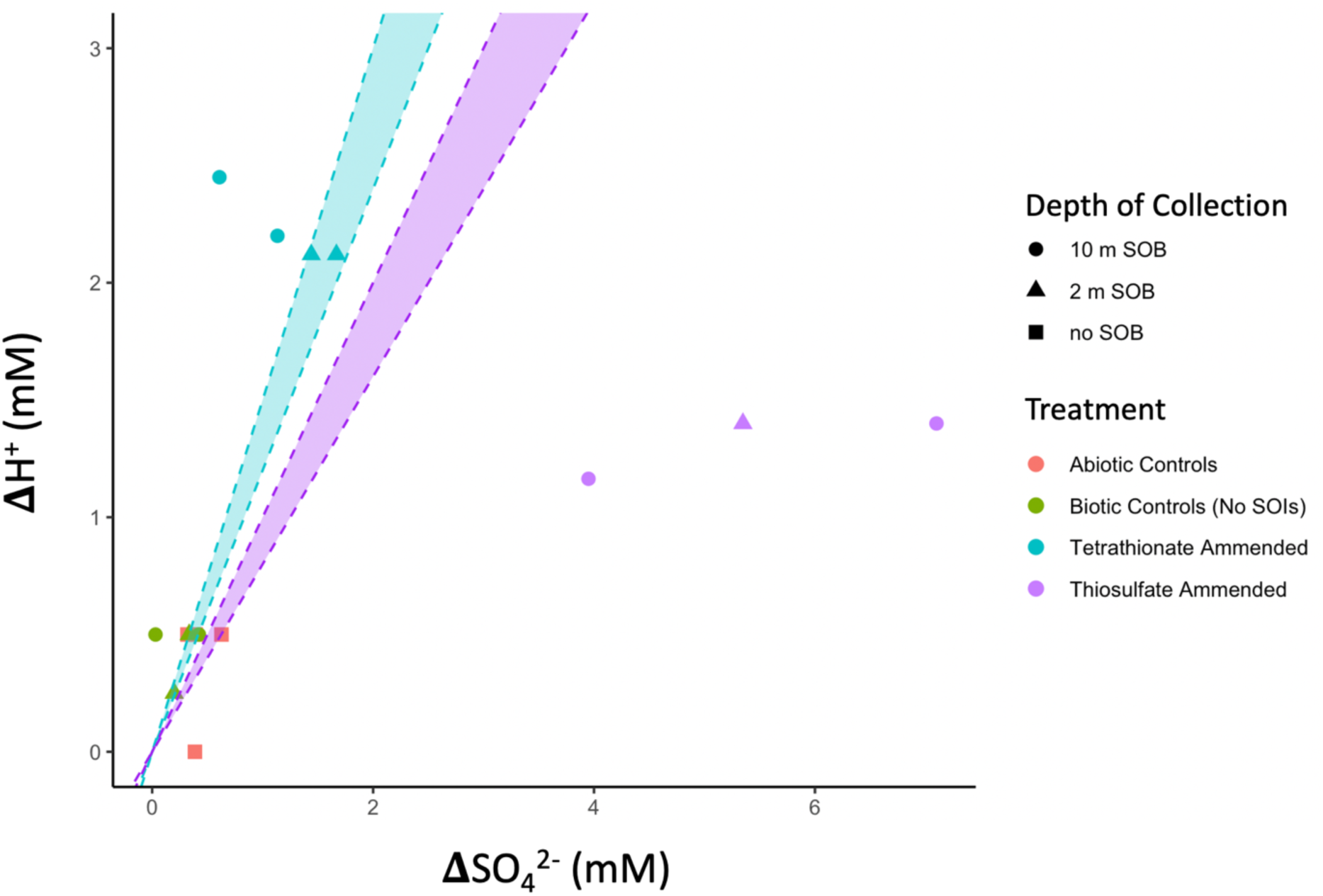
The actual ΔH^+^ : ΔSO_4_^2−^ ratio across both tetrathionate- (blue dots) and thiosulfate- (purple dots) amended experiments fall below the 3:2 theoretical ratio predicted from direct tetrathionate oxidation to sulfate (blue dashed lines and zone represent the ratio determined from the following equation, with an ion activity constant range of 0.8-1.0 for H^+^ according to S_4_O_6_^2−^ + 3.5 O_2_ + 3 H_2_O → 4 SO_4_^2−^ + 6 H^+^) or the 1:1 ratio predicted for direct thiosulfate oxidation to sulfate via cSOx (purple dashed line zone represent the ratio determined from the following equation, with an ion activity constant range of 0.8-1.0 for H^+^ according to cSOx: S_2_O_3_^2−^ + H_2_O + 2 O_2_ → 2 SO_4_^2−^ + 2 H^+^). Actual ΔH^+^ for SOB- and SOI-amended systems was calculated as (Δ[H^+^] = [H^+^] at t_end_ – [H^+^] at t_0_ + H^+^ consumed by buffer, as calculated using VMinteq, see methods for calculation details).

The proportion of genera in these communities shifted over the course of the experiments. At t_0_, the inocula for experiments A and B (both 2 and 10 m communities) were dominated by *Pandoraea* (12–74% of the total microbial community), while *Halothiobacillus* (<1%) and *Thiomonas* (<1%) were also present. For experiment C, the 2 m and 10 m inoculum were dominated by *Halothiobacillus* (94–97%), while *Thiomonas* (2–3%) and *Pandoraea* (<1%) were also present (Fig 4, SI Table 4). By t_end_, the SOB communities became (for Experiments A and B) or remained (for Experiment C) dominant in all but one of the SOI-amended microcosms (median proportion of SOB in SOI treated t_end_ = 87%, ^F^i^g^ 4a), indicating that SOI compounds were an energy source. The remaining microcosm, where SOB genera at t_end_ = 7%, instead contained a high abundance of *Pseudomonas*, explaining the point of inflection in pH, switching from acidity-generating to - consuming that occurred only in this system (SI Table 3). In contrast to microcosms which received sulfur substrate, where the abundance of SOB was high, only two of the microcosms which did not receive sulfur substrate contained a large fraction of *Halothiobacillus* at t_end_ (78–89%, Fig 4); in the other two SOI-free treatments, *Halothiobacillus* concentrations at t_end_ decreased to <1% (with no pattern with depth or substrate, Fig 4). In these microcosms without sulfur substrate, *Pandoraea* — the most abundant SOB — only comprised 9–12% of the total microbial community. Most of the remaining community in these biotic controls was composed of the heterotrophic genera *Delftia* and *Pseudomonas* (SI Fig 2, SI Table 4).

**Fig 5.**
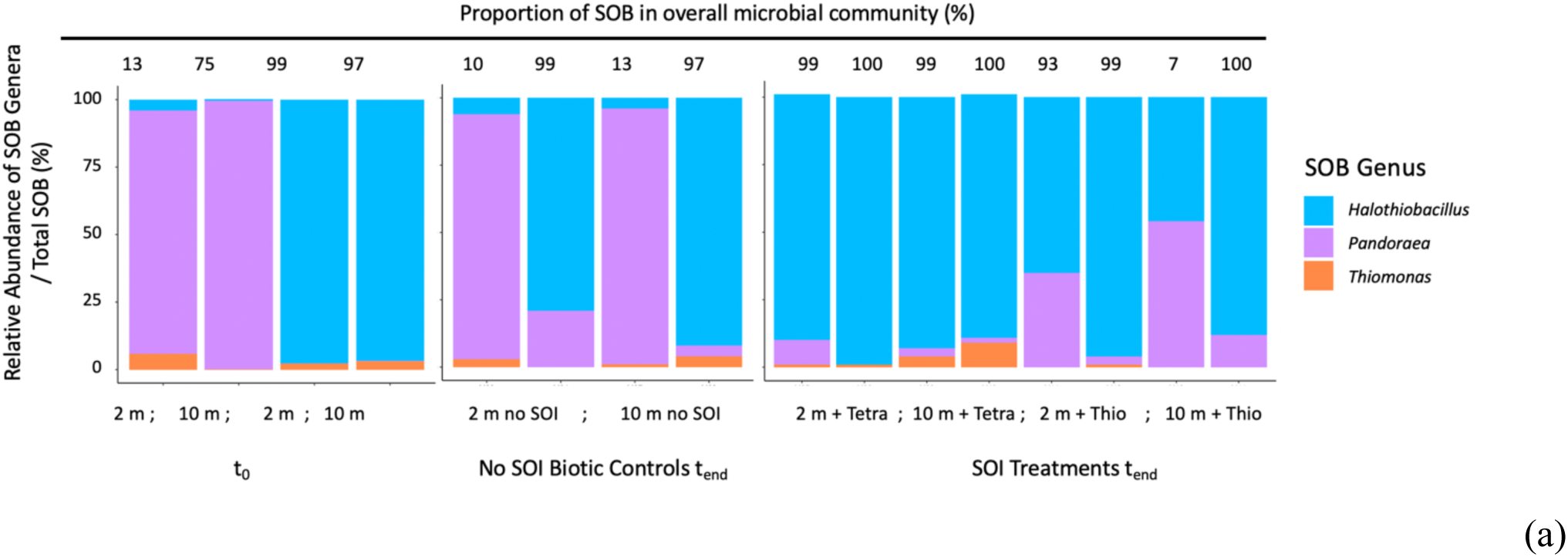
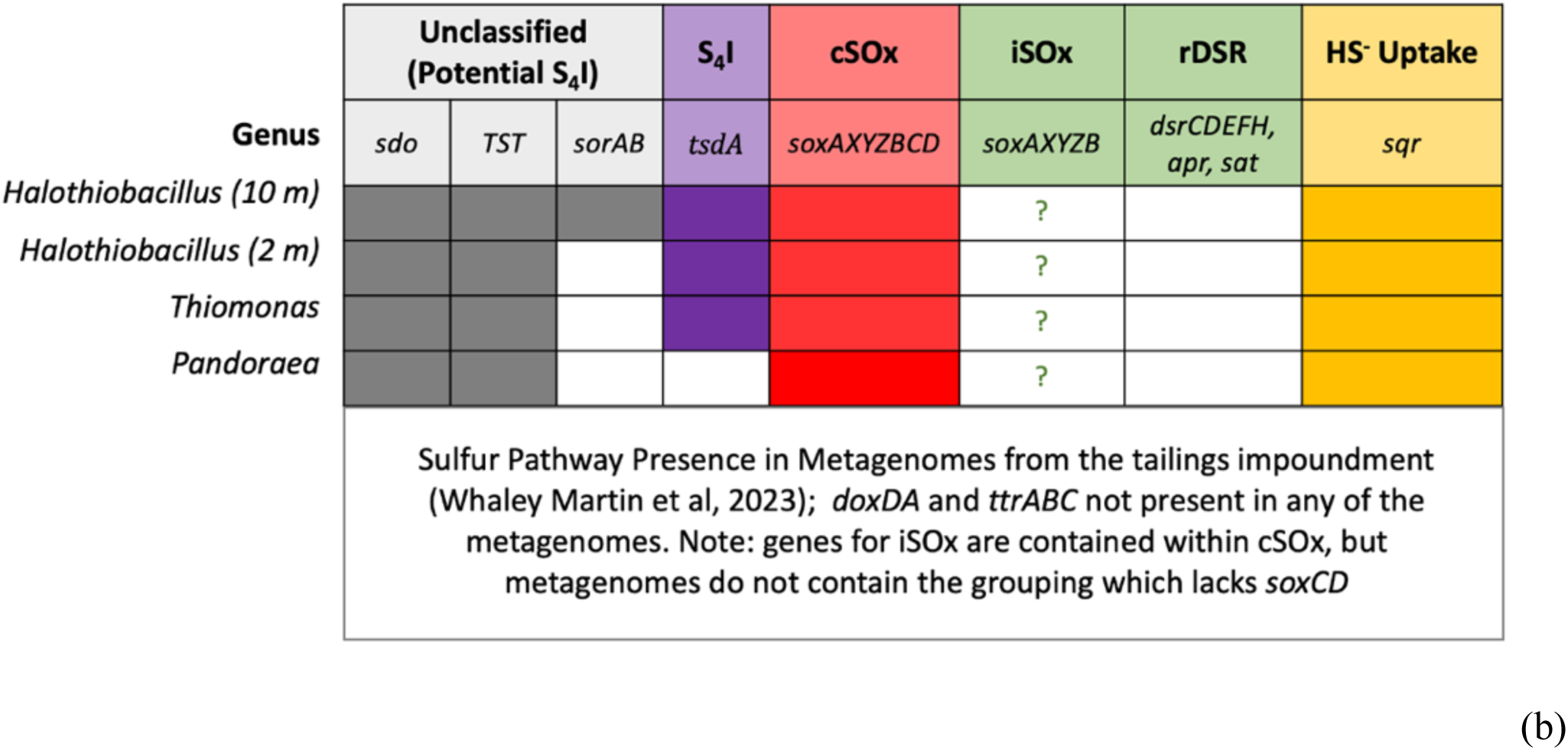
(a) The SOB community was composed of *Halothiobacillus*, *Pandoraea*, and *Thiomonas,* isolated from samples taken from either 2 m or 10 m depths of the tailings impoundment, and forming the bulk of the communities at t_0_ and t_end_. Regardless of whether *Halothiobacillus* was dominant in the initial communities, the genus comprised the largest proportion of the SOB population in microcosms that received amendments of thiosulfate or tetrathionate, although *Pandoraea* comprised a large fraction in one of the thiosulfate-amended microcosms. (b) Metagenomic data for *Halothiobacillus*, *Thiomonas* and *Pandoraea* communities indicates that all three genera contain genes for the cSOx pathway (*soxAX, soxYZ, soxCD* and *soxB*), and therefore contain the genes required for iSOx (*soxAX, soxYZ,* and *soxB*). However, the iSOx pathway formed of *soxAX, soxYZ,* and *soxB* but lacking *soxCD* was not detected. *Halothiobacillus* and *Thiomonas* genera also contain the first stage of the S_4_I pathway (*tsdA*), although *Pandoraea* did not. They also contained genes for sulfide oxidation (*sqr*), and potentially the oxidation of zero valence sulfur (ZVS) to sulfite (*sdo*). Metagenomes from 10 m communities indicate they have the ability to form sulfate from sulfite (*sor*), see SI Table 4.

Metagenomes from both *Halothiobacillus* and *Thiomonas* (originally sequenced by Whaley-Martin et al, 2023) indicate that these genera can process thiosulfate via two pathways: the cSOx pathway (*soxAX*, *soxYZ*, *soxCD*, and *soxB*) or the first stage of the S_4_I pathway (*tsdA*, Fig 4). While *Pandoraea* metagenomes from this study site were not available, metagenomes sequenced from a different mine site, combined with a review of 64 metagenomes, indicate that this genus contains cSOx but not TsdA (Fig 4b). In addition, both *Pandoraea* and *Halothiobacillus* from the 2 m and 10 m communities also contain the genes for *sdo* (S^0^ → SO_3_^2−^), although only the 10 m community contained *sor* (SO_3_^2−^ → SO_4_^2−^). It is also worth noting that *Halothiobacillus* and *Pandoraea* cells can reduce nitrite to ammonia (*nirB*) but lack the capacity to reduce nitrate to nitrite (*narGHIJ* / *napAB*, see SI Table 5). Bacteria belonging to the genus *Thiomonas* have the complimentary capacity to reduce nitrate to nitrite (*narGHIJ*) but lack the genes for nitrite reductase (*nirB, nirSK*).

### Dissolved Oxygen and Nitrate as Terminal Electron Acceptors

During the oxic experiments (Experiments A and B, and the first 28–30 hours of Experiment C), oxygen concentrations decreased rapidly in all SOB inoculated microcosms, but not in the abiotic controls (Table 2, SI Table 3). Although a greater concentration of oxygen was consumed by the heterotrophic genera *Delftia* and *Pseudomonas* without sulfur substrate (SI Fig 3, SI Table 3), this observation supports the prediction that oxygen is the terminal electron acceptor (TEA) for sulfur oxidation when available.

However, no obvious TEA was paired with thiosulfate oxidation under suboxic conditions. Although established under oxic conditions, the thiosulfate-amended microcosms in Experiment C became anoxic by ∼ 30 h (SI Fig 3, SI Table 3). Additional tests to examine the oxygen supply via surface diffusion indicated that SOB microcosms were anoxic at all probe depths, and ingress of oxygen via surface diffusion was insubstantial over the course of the incubation period (0.75 µM/h). During the period of suboxia, ([DO] < 1 μM) there was no statistically significant change in the concentrations of nitrogen species (nitrate, nitrite, ammonia; SI Table 3). Further, the only SOB with the capacity to reduce nitrate, *Thiomonas*, was found at very low relative abundances in the suboxic experiments (Fig 4). Both lines of evidence indicate that the low proton yield cannot be accounted for by a shift to thiosulfate oxidation using nitrate. Since no other oxidants, such as manganese or iron, were readily available in the medium (2.0 mM MgSO_4_, 0.2 mM NH_4_Cl, 1.4 mM K_2_HPO_4_, 8.3 mM NaNO_3_, and trace elements), this suggests thiosulfate was metabolised by SOB in these microcosms with an atypical electron acceptor.

## Discussion

### Sulfur Reactions the Oxic Tetrathionate-Amended Microcosms

The production of acid in the oxic microcosms amended with tetrathionate, indicates that latter stage S_4_I pathway reactions were active in *Halothiobacillus* and *Pandoraea* dominated SOB communities. These genera either oxidized tetrathionate (a) directly to sulfate or (b) indirectly to other polythionates, and perhaps even ZVS, some of which subsequently oxidized to sulfate. In these microcosms, the low ΔH^+^ : ΔSO_4_^2−^ yield was likely a result of phosphate buffering in the media, so does not disproved direct oxidation. However, the small concentrations of thiosulfate formed and consumed are evidence suggesting that cycling through higher order polythionate pools may have occurred.

Members of the genus *Halothiobacillus*, which grew to compose the majority of the tetrathionate-amended communities, are also observed to process tetrathionate in other contexts. When isolated strains of *H. neapolitanus* were cultured under oxic conditions, they thrived on both thiosulfate or tetrathionate as energy sources (Wood et al. 2005). *H. kellyi*, isolated from hydrothermal vents in the Aegean sea, was also found to be capable of growth on thiosulfate and tetrathionate under oxic conditions (Sievert et al. 2000).

However, known sulfur enzyme mechanisms are insufficient to explain the changes in sulfur speciation observed in these microcosms. Direct oxidation of tetrathionate to sulfate appears the most likely process. However, the enzymes thought to compose the latter half of the S_4_I pathway (equation 2) fall broadly into two mechanisms, neither of which is a simple direct oxidation. Some mechanisms, such as those proposed by Pyne et al. (SoxB and SoxCD), are believed to split tetrathionate into sulfate and ZVS (Pyne et al. 2018). Although SoxB is present in the SOB metagenomes, the mechanism does not explain the generation of thiosulfate during Experiments A and B. However, the majority of proposed mechanisms, such as those facilitated by TetH, are thought to disproportionate tetrathionate into thiosulfate, sulfate and ZVS, possibly via a process that generates higher-order SOIs, such as trithionate [S_3_O_6_^2−^, SI Table 2, (Kanao et al. 2007; Beard et al. 2011; Rameez et al. 2020; Zhang et al. 2020; Cai et al. 2022)]. The TetH pathway is described in *Acidothiobacillus caldus* and *A. ferrooxidans* as a reaction that forms disulfane monosulfonic acid (DSMSA), which then spontaneous decomposes to thiosulfate and ZVS, as shown in equation 4 (De Jong et al. 1997; Dahl 2005; Kanao et al. 2007; Beard et al. 2011; Yu et al. 2014; Wang et al. 2019; Camacho et al. 2020b; Gwak et al. 2022):

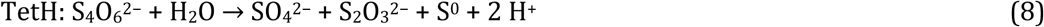

While there is consensus that TetH results in the formation of sulfate, thiosulfate, and ZVS, Beard et al. proposed that the intermediary, DSMSA, is also capable of forming other polythionates (S_6_O_6_^2−^, S_5_O_6_^2−^) under both aerobic and anaerobic conditions, opening the door to polythionate cycling (Beard et al. 2011). Pentathionate has also occasionally been detected as a product of tetrathionate hydrolase, and may be a polythionate formed in these microcosms (Bugaytsova and Lindström 2004). *Thiomonas intermedia K12* oxidation of tetrathionate was previously shown to result in thiosulfate, sulfate, and trithionate (speculated to form from DSMSA following the hydrolysis of tetrathionate), suggesting that trithionate may have also formed in this experiment (Wentzien and Sand 2004). However, in these microcosms, only low concentrations of thiosulfate and S^0^ were detected, reducing support for the TetH mechanism. This suggests an alternate enzyme, yet to be identified, exists which facilitates direct tetrathionate oxidation.

### Sulfur Reactions the Oxic Thiosulfate-Amended Microcosms

Over this oxic period in Experiment C (<30 h, changes calculated from 0–24 hours), both the ΔH^+^ : ΔSO_4_^2−^ yield and the ΔSO_4_^2−^ : ΔS-S_2_O_3_^2−^ ratio are within the theoretical range accounted for by the cSOx pathway was the sole sulfur oxidation reaction, once phosphate buffering in the system was accounted for. Parallel activation of the cSOx and S_4_I Part 1 pathways might also contribute to the low proton yield, as was previously proposed to explain the low ΔH^+^ : ΔSO_4_^2−^ ratios observed in 500 L mesocosm experiments in 2020 (Gordon et al. 2024).

### Sulfur Reactions the Suboxic Thiosulfate-Amended Microcosms

Previous research identified the key role the genera *Halothiobacillus* and *Thiomonas* in producing acidity in mine wastewater systems via the cSOx pathway (Whaley-Martin et al. 2019; Gordon et al. 2024a). As a result of this study, *Pandoraea* should be added to this suite of cSOx guild SOB in mine wastewaters. The proton yield (ΔH^+^: ΔSO_4_^2−^) remained extremely low during the suboxic experimental duration, although it was slightly higher than during the oxic portion. Similarly, low ΔH^+^ : ΔSO_4_^2−^ ratios have been observed previously for *Halothiobacillus* isolated from this wastewater systems, where the calculated ratio was 0–0.3 (Whaley-Martin et al. 2019), although phosphate buffering would also have been present in this media (1.1% (w/v) K_2_HPO_4_). While low proton yields might, again, be accounted for by phosphate buffering, the sulfur speciation data clearly indicate that the cSOx pathway was not the sole active mechanism for metabolism in these microcosms. Although 4.2 mM of thiosulfate was converted to sulfate over 118 h, and the remaining 7.5 mM was converted to unknown aqueous sulfur species based on the sulfur mass balance (a comparison of TotS at t_0_ and t_end_ indicates that S was not lost as H_2_S_(g)_). This indicates formation of unmeasured sulfur species, because the small concentration (<1.5 mM) of S^0^ formed at the midpoint of the experiments was consumed by t_end_.

Further, during the suboxic portion of the experiment (deltas calculated from 74–192 h), a lack of e^−^ acceptors was observed as thiosulfate concentrations decreased by ∼12 mM. During this period, no DO was detected at any probe depth, and surface diffusion would account for <1 µM/h. Although nitrate was present at sufficient concentrations for thiosulfate oxidation, neither [NO_3_^−^] nor [NO_2_^−^] decreased to a degree that could account for their use as TEA, and the population of *Thiomonas* (the only genus with capacity to reduce NO_3_^−^) was negligible in these microcosms. In contrast, mine wastewater systems sometimes contain populations of *Thiobacillus*, whose use of nitrate as an electron acceptor has been previously demonstrated (Chapter 4; Whaley Martin et al, 2023).

Thiosulfate disproportionation, which might a potential mechanism for thiosulfate metabolism without an e^−^ acceptor, is not supported by these experiments. Thiosulfate disproportionation according to equation 9 (Jackson and McInerney 2000) can be eliminated as a possible mechanism, as it would produce HS^−^/H_2_S_(g)_ (1:1 ratio of HS^−^ to H_2_S_(g)_ would be expected at pH 7; below this pH, H_2_S_(g)_ rapidly dominates, reaching 100% H_2_S_(g)_ by pH 5.5) and no known enzyme system can catalyze the reaction:

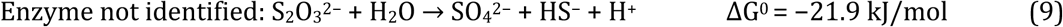

Similarly, abiotic thiosulfate disproportionation is only spontaneous below pH 4, and therefore can be ruled out (Gwak et al. 2022). Finally, hydrolysis of thiosulfate via iSOx (contained within the cSOx genes), forms similar products to those formed via disproportionation (equation 10):

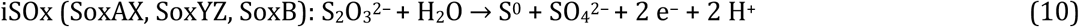

However, although the shift between cSOx and iSOx pathways has been observed in *Sulfurimonas* (Wang et al. 2021), there is no evidence of iSOx use by *Halothiobacillus*, *Thiomonas*, or *Pandoraea*. Further, high concentrations of the product S^0^ were not observed in experiment C, and if iSOx were employed, an e^−^ acceptor would still be required.

The survival of SOB under suboxic conditions, indicated by the high proportion of *Halothiobacillus* and *Pandoraea* at t_end_ in the treatments amended with thiosulfate (visual observations indicated increased cell density over the course of the experiment) is particularly intriguing, as the literature reports that these genera are strictly aerobic. Although strictly aerobic, *Halothiobacillus* has occasionally been found to oxidize sulfur under very low oxygen conditions (Magnuson et al. 2023). If the efficiency of S_4_I pathway enzymes were to fall between the efficiency calculated for a genus known to use cSOx (*Halothiobacillus neapolitanus*, E = 10%) and one that uses rDSR (*Thiobacillus denitrificans*, E = 22%), pathway would be advantageous under low oxygen conditions (Klatt and Polerecky 2015).

Limited oxygen use by *Halothiobacillus* under suboxic conditions could be a potential explanation for the observations made in these experiments, particularly near the surface of the microcosms. An approximation of oxygen utilization rates by *Halothiobacillus*-dominated biofilms, based on biofilm oxygen penetration depths (Satoh et al. 2009) and oxygen utilization rates with sulfur oxidation observed in batch reactors (Jensen et al. 2011) suggest that biotic oxygen consumption might increase to 4 µM/h or a total of 0.7 mM over the anoxic portion of the experiment (SI Table 6). Activation of the S_4_I pathway P1 would be a plausible mechanism for reducing acidity generation, and it might be possible under microaerophilic conditions if SOB increased surface oxygen draw to ∼1 mM over the anoxic potion of the experiment (equations 11–12). This would produce an experimental ratio of ∼12 ΔS-S_2_O_3_^2−^ : 1 ΔO_2_, where the theoretical ratio is 8 ΔS-S_2_O_3_^2−^ : 1 ΔO_2_ (equations 11–12):

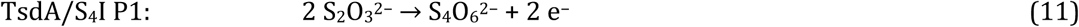

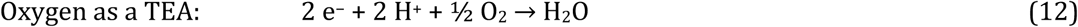

Near the surface, *Halothiobacillus* in the thiosulfate-amended microcosms may be surviving on the low concentrations of oxygen in the water introduced through surface diffusion. However, even if low concentrations of oxygen were available at the surface of the microcosm, thiosulfate diffusion rates suggest that molecules near the bottom of the flask would not have time to come into contact with the surface layer (SI Table 7); therefore, an alternate e^−^ acceptor must have been employed. In addition, activity of the first stage of the S_4_I pathway would not account for sulfate formation during this period. Manganese, the e^−^ acceptor whose reduction is next most favorable on the redox ladder, was not present in the media (see 5.2.1). With no other possible e^−^ acceptor, polythionate reduction at the bottom of the 6 L microcosms may have been paired with thiosulfate oxidation to sulfate under suboxic conditions. Polythionates have been previously observed in SOB communities of *Halothiobacillus*, *Thiomonas*, and *Pandoraea*. Under anaerobic conditions, other studies observed sulfur, sulfite, and trithionate formation with thiosulfate oxidation in *Pandoraea*, as well as a small amount of trithionate formation with thiosulfate loss in *Halothiobacillus* (Anandham et al. 2008). *Thiomonas intermedia K12* (isolated from sewage) was observed to degrade tetrathionate to produce primarily pentathionate and small amounts of hexathionate, thiosulfate, trithionate, and sulfate (Wentzien and Sand 2004). Sequencing mRNA may provide further insights into the sulfur enzymes active in the system.

### Relevance to Mine Wastewater Systems

These microcosms demonstrate that proton yields resulting from thiosulfate and tetrathionate oxidation by SOB fall far below those theoretically predicted. Tetrathionate oxidation produces nearly the ΔH^+^: ΔSO_4_^2−^ theoretically predicted, but thiosulfate oxidation produces < 1/2. Given these ratios, is sulfur-driven acid generation a concern to mine wastewater systems?

Potential impacts can vary greatly considering typical SOI concentrations across four Canadian mine tailings reservoirs sampled in 2018: thiosulfate (S–S_2_O_3_^2−^) < 0.003–0.8 mM, trithionate (S–S_3_O_6_^2−^) < 0.03–0.1 mM, and tetrathionate (S–S_4_O_6_^2−^) < 0.03–0.23 mM (K. Whaley-Martin et al. 2020). If 0.2 mM S–S_2_O_3_^2−^ oxidation were to produce acid according to the ratio in observed in the anoxic microcosms (0.02:1 H^+^/ S–S_2_O_3_^2−^), this would produce 0.004 mM H^+^, which would decrease the pH from 7 to 5.4. If the same concentration of tetrathionate (0.2 mM S–S_4_O_6_^2−^) was to produce the acid ratio observed in the tetrathionate-amended microcosms (ΔH^+^ : ΔSO_4_^2−^ [assumed from S-S_4_O_6_^2−^] ∼ 2.2:1), this would produce 0.44 mM H^+^, which would decrease the pH from 7 to 3.4. However, buffering in the water would reduce these impacts, perhaps even neutralize them completely. With this range in potential outcomes from SOI concentrations observed at active mine sites, we need greater clarity as to which SOB-facilitated SOI reactions occur in these systems; this continues to be an important future research direction.

## Conclusions

In mine wastewaters, the activities of sulfur pathways used by *Halothiobacillus, Thiomonas*, and *Pandoraea* are important, as they impact the ability to predict and prevent acidity generation in tailings impoundments. These experiments demonstrate that acidity generation will occur immediately with both tetrathionate oxidation under oxic conditions and with thiosulfate oxidation under oxic and anoxic conditions, yet at ratios much lower than would theoretically be predicted. The low initial acid generation, in combination with gaps in the sulfur mass balance, indicate that S redox processes occur that are not currently linked to known sulfur enzyme systems. It is possible that polythionates are forming which add complexity to the tetrathionate oxidation, and further studies are needed to investigate which enzyme-facilitated reactions occur within this polythionate pool. To date, research into this pool is limited as standards for these compounds (S_3_O_6_^2−^, S_4_O_6_^2−^, S_5_O_6_^2−^) are not readily available and robust analytical techniques need to be developed. In addition, these SOB communities demonstrated an ability to oxidize thiosulfate under suboxic conditions with no apparent TEA, which is a phenomenon requiring further investigation.

These observations contrast with the previous observations that *Halothiobacillus*-dominated SOB communities amended with thiosulfate always produce acidity through cSOx pathway activation (Whaley-Martin et al. 2023a). Therefore, the interplay between cSOx and S_4_I pathway in genera such as *Halothiobacillus* is an important consideration when predicting mine wastewater acidity. For applications in mine wastewater management, 16S rRNA gene sequences are a technique which can be immediately applied to screen for SOB communities. However, because of the presence of multiple pathways in several SOB genera, there is additional value in developing targeted screening for the activation of specific enzymes such as SoxCD, TsdA, TetH, and DsrC. Further research is also required to clarify the specifics of polythionate cycling in mine wastewater systems. Finally, the development of treatments to promote the use of the cSOx pathway by the SOB community would promote in immediate acid generation, allowing onsite treatment.

## Acknowledgements

The author would like to thank on-site mine personnel who aided in site orientation and collection of samples from which SOB communities were isolated. Thanks go to Dr. Kelly Whaley-Martin who cryopreserved the 2017 SOB cultures, isolated from an active tailings reservoir, used as inocula for these experiments. Further, I would like to thank Dr. Dalal Askar for equipment training and maintenance, analytical method improvements, and research guidance. I would also like to thank Lauren Twible for demonstrating correct techniques for 16S analysis, ensuring QA/QC. In addition, I am very grateful to the efforts of Joshua Crawford and In Him Lee, who assisted in setting up microcosms, collecting samples, performing aqueous geochemical analyses and DNA extractions. In Him Lee also developed python script for consolidating and analysing pH and DO probe data, which aided in data analysis. Finally, many thanks go out to Dr. Simon Apte (S.A.) who provided coaching and guidance through the drafting of the manuscript and Dr. Lesley Warren (L.A.W.) for funding/facilities.

## References

Anandham R, Indiragandhi P, Kwon SW, et al (2010) Pandoraea thiooxydans sp. nov., a facultatively chemolithotrophic, thiosulfate-oxidizing bacterium isolated from rhizosphere soils of sesame (Sesamum indicum L.). Int J Syst Evol Microbiol 60:21–26. 10.1099/ijs.0.012823-0

Anandham R, Indiragandhi P, Madhaiyan M, et al (2008) Chemolithoautotrophic oxidation of thiosulfate and phylogenetic distribution of sulfur oxidation gene (soxB) in rhizobacteria isolated from crop plants. Res Microbiol 159:579–589. 10.1016/j.resmic.2008.08.007

Anantharaman K, Hausmann B, Jungbluth SP, et al (2018) Expanded diversity of microbial groups that shape the dissimilatory sulfur cycle. ISME J 12:1715–1728. 10.1038/s41396-018-0078-0

Arsène-Ploetze F, Koechler S, Marchal M, et al (2010) Structure, Function, and Evolution of the Thiomonas spp. Genome. PLoS Genet 6:e1000859. 10.1371/journal.pgen.1000859

Beard S, Paradela A, Albar JP, Jerez CA (2011) Growth of Acidithiobacillus Ferrooxidans ATCC 23270 in Thiosulfate Under Oxygen-Limiting Conditions Generates Extracellular Sulfur Globules by Means of a Secreted Tetrathionate Hydrolase. Front Microbiol 2:. 10.3389/fmicb.2011.00079

Bernier L, Warren LA (2007) Geochemical diversity in S processes mediated by culture-adapted and environmental-enrichments of Acidithiobacillus spp. Geochim Cosmochim Acta 71:5684–5697. 10.1016/j.gca.2007.08.010

Brito JA, Denkmann K, Pereira IAC, et al (2015) Thiosulfate Dehydrogenase (TsdA) from Allochromatium vinosum. J Biol Chem 290:9222–9238. 10.1074/jbc.M114.623397

Bugaytsova Z, Lindström EB (2004) Localization, purification and properties of a tetrathionate hydrolase from Acidithiobacillus caldus. Eur J Biochem 271:272–280. 10.1046/j.1432-1033.2003.03926.x

Cai R, He W, Liu R, et al (2022) Deep-Sea In Situ Insights into the Formation of Zero-Valent Sulfur Driven by a Bacterial Thiosulfate Oxidation Pathway. MBio 13:. 10.1128/mbio.00143-22

Camacho D, Frazao R, Fouillen A, et al (2020a) New Insights Into Acidithiobacillus thiooxidans Sulfur Metabolism Through Coupled Gene Expression, Solution Chemistry, Microscopy, and Spectroscopy Analyses. Front Microbiol 11:. 10.3389/fmicb.2020.00411

Camacho D, Jessen GL, Mori JF, et al (2020b) Microbial Succession Signals the Initiation of Acidification in Mining Wastewaters. Mine Water Environ 39:669–683. 10.1007/s10230-020-00711-9

Camacho D, Jessen GL, Mori JF, et al (2020c) Microbial Succession Signals the Initiation of Acidification in Mining Wastewaters. Mine Water Environ 39:669–683. 10.1007/s10230-020-00711-9

Chen L, Hu M, Huang L, et al (2015) Comparative metagenomic and metatranscriptomic analyses of microbial communities in acid mine drainage. ISME J 9:1579–1592. 10.1038/ismej.2014.245

Chen X-G, Geng A-L, Yan R, et al (2004) Isolation and characterization of sulphur-oxidizing Thiomonas sp. and its potential application in biological deodorization. Lett Appl Microbiol 39:495–503. 10.1111/j.1472-765X.2004.01615.x

Chua K-O, See-Too W-S, Ee R, et al (2019) In silico Analysis Reveals Distribution of Quorum Sensing Genes and Consistent Presence of LuxR Solos in the Pandoraea Species. Front Microbiol 10:. 10.3389/fmicb.2019.01758

Dahl C (2005) Environmental Technologies to Treat Sulfur Pollution - Principles and Applications, 2nd Editio. IWA Publishing

De Jong GAH, Hazeu W, Bos P, Kuenen JG (1997) Isolation of the tetrathionate hydrolase from Thiobacillus acidophilus. Eur J Biochem 243:678–683. 10.1111/j.1432-1033.1997.00678.x

Denef VJ, Mueller RS, Banfield JF (2010) AMD biofilms: Using model communities to study microbial evolution and ecological complexity in nature. ISME J 4:599–610. 10.1038/ismej.2009.158

Denkmann K, Grein F, Zigann R, et al (2012) Thiosulfate dehydrogenase: a widespread unusual acidophilic c -type cytochrome. Environ Microbiol 14:2673–2688. 10.1111/j.1462-2920.2012.02820.x

Du R, Gao D, Wang Y, et al (2022) Heterotrophic Sulfur Oxidation of Halomonas titanicae SOB56 and Its Habitat Adaptation to the Hydrothermal Environment. Front Microbiol 13:. 10.3389/fmicb.2022.888833

Eckley CS, Luxton TP, McKernan JL, et al (2015) Influence of reservoir water level fluctuations on sediment methylmercury concentrations downstream of the historical Black Butte mercury mine, OR. Appl Geochemistry 61:284–293. 10.1016/j.apgeochem.2015.06.011

Friedrich CG, Rother D, Bardischewsky F, et al (2001) Oxidation of Reduced Inorganic Sulfur Compounds by Bacteria: Emergence of a Common Mechanism? Appl Environ Microbiol 67:2873–2882. 10.1128/AEM.67.7.2873-2882.2001

Ghosh W, Dam B (2009) Biochemistry and molecular biology of lithotrophic sulfur oxidation by taxonomically and ecologically diverse bacteria and archaea. FEMS Microbiol Rev 33:999–1043. 10.1111/j.1574-6976.2009.00187.x

Gordon J, Apte SC, Colenbrander Nelson TE, et al (2024) Microbial Sulfur Pathways and Outcomes in Tailings Impoundments: A Mesocosm Study. Mine Water Environ. 10.1007/s10230-024-01016-x

Grettenberger CL, Havig JR, Hamilton TL (2019) Metabolic diversity and co-occurrence of multiple Ferrovum species at an acid mine drainage site. bioRxiv 1–14. 10.1101/751859

Gwak JH, Awala SI, Nguyen NL, et al (2022) Sulfur and methane oxidation by a single microorganism. Proc Natl Acad Sci U S A 119:1–12. 10.1073/pnas.2114799119

Haja DK, Wu C-H, Poole FL, et al (2020) Characterization of thiosulfate reductase from Pyrobaculum aerophilum heterologously produced in Pyrococcus furiosus. Extremophiles 24:53–62. 10.1007/s00792-019-01112-9

Han Y, Perner M (2015) The globally widespread genus Sulfurimonas: Versatile energy metabolisms and adaptations to redox clines. Front Microbiol 6:1–17. 10.3389/fmicb.2015.00989

Hao T wei, Xiang P yu, Mackey HR, et al (2014) A review of biological sulfate conversions in wastewater treatment. Water Res 65:1–21. 10.1016/j.watres.2014.06.043

Heinzinger NK, Fujimoto SY, Clark MA, et al (1995) Sequence analysis of the phs operon in Salmonella typhimurium and the contribution of thiosulfate reduction to anaerobic energy metabolism. J Bacteriol 177:2813–2820. 10.1128/jb.177.10.2813-2820.1995

Hensel M, Hinsley AP, Nikolaus T, et al (1999) The genetic basis of tetrathionate respiration in Salmonella typhimurium. Mol Microbiol 32:275–287. 10.1046/j.1365-2958.1999.01345.x

Hinsley AP, Berks BC (2002) Specificity of respiratory pathways involved in the reduction of sulfur compounds by Salmonella enterica. Microbiology 148:3631–3638. 10.1099/00221287-148-11-3631

Hua Z-S, Han Y-J, Chen L-X, et al (2015) Ecological roles of dominant and rare prokaryotes in acid mine drainage revealed by metagenomics and metatranscriptomics. ISME J 9:1280–1294. 10.1038/ismej.2014.212

Jackson BE, McInerney MJ (2000) Thiosulfate disproportionation by Desulfotomaculum thermobenzoicum. Appl Environ Microbiol 66:3650–3653. 10.1128/AEM.66.8.3650-3653.2000

Jenner LP, Kurth JM, Helmont S Van, et al (2019) Heme ligation and redox chemistry in two bacterial thiosulfate dehydrogenase (TsdA) enzymes. J Biol Chem 294:18002–18014. 10.1074/jbc.RA119.010084

Jensen HS, Lens PNL, Nielsen JL, et al (2011) Growth kinetics of hydrogen sulfide oxidizing bacteria in corroded concrete from sewers. J Hazard Mater 189:685–691. 10.1016/j.jhazmat.2011.03.005

Kanao T, Kamimura K, Sugio T (2007) Identification of a gene encoding a tetrathionate hydrolase in Acidithiobacillus ferrooxidans. J Biotechnol 132:16–22. 10.1016/j.jbiotec.2007.08.030

Kappler U, Friedrich CG, Trüper HG, Dahl C (2001) Evidence for two pathways of thiosulfate oxidation in Starkeya novella (formerly Thiobacillus novellus). Arch Microbiol 175:102–111. 10.1007/s002030000241

Klatt JM, Polerecky L (2015) Assessment of the stoichiometry and efficiency of CO2 fixation coupled to reduced sulfur oxidation. Front Microbiol 6:484. 10.3389/fmicb.2015.00484

Kletzin A (1992) Molecular characterization of the sor gene, which encodes the sulfur oxygenase/reductase of the thermoacidophilic archaeum Desulfurolobus ambivalens. J Bacteriol 174:5854–5859. 10.1128/jb.174.18.5854-5859.1992

Li W, Zhang M, Kang D, et al (2020) Mechanisms of sulfur selection and sulfur secretion in a biological sulfide removal (BISURE) system. Environ Int 137:105549. 10.1016/j.envint.2020.105549

Lopes AR, Madureira D, Diaz A, et al (2020) Characterisation of bacterial communities from an active mining site and assessment of its potential metal solubilising activity. J Environ Chem Eng 8:104495. 10.1016/j.jece.2020.104495

Lu W-P, Kelly DP (1988) Cellular Location and Partial Purification of the “Thiosulphate-oxidizing Enzyme” and “Trithionate Hydrolyase” from Thiobacillus tepidarius. Microbiology 134:877– 885. 10.1099/00221287-134-4-877

Magnuson E, Altshuler I, Freyria NJ, et al (2023) Sulfur-cycling chemolithoautotrophic microbial community dominates a cold, anoxic, hypersaline Arctic spring. Microbiome 1–20. 10.1186/s40168-023-01628-5

Meulenberg R, Pronk JT, Frank J, et al (1992a) Purification and partial characterization of a thermostable trithionate hydrolase from the acidophilic sulphur oxidizer Thiobacillus acidophilus. Eur J Biochem 209:367–374. 10.1111/j.1432-1033.1992.tb17298.x

Meulenberg R, Pronk JT, Hazeu W, et al (1992b) Oxidation of reduced sulphur compounds by intact cells of Thiobacillus acidophilus. Arch Microbiol 157:161–168. 10.1007/BF00245285

Miettinen H, Bomberg M, Le TMK, Kinnunen P (2021) Identification and metabolism of naturally prevailing microorganisms in zinc and copper mineral processing. Minerals 11:1–31. 10.3390/min11020156

Miranda-Trevino JC, Pappoe M, Hawboldt K, Bottaro C (2013) The Importance of Thiosalts Speciation: Review of Analytical Methods, Kinetics, and Treatment. Crit Rev Environ Sci Technol 43:2013–2070. 10.1080/10643389.2012.672047

Müller FH, Bandeiras TM, Urich T, et al (2004) Coupling of the pathway of sulphur oxidation to dioxygen reduction: Characterization of a novel membrane-bound thiosulphate:quinone oxidoreductase. Mol Microbiol 53:1147–1160. 10.1111/j.1365-2958.2004.04193.x

Musuku B, Kasymova D, Saari E, Dahl O (2023) Influence of Water Quality on Sulphide Ore Oxidation and Speciation of Sulphur Anions during Autogenous Milling. Minerals 13:277. 10.3390/min13020277

Opara CB, Kamariah N, Spooren J, et al (2023) Interesting Halophilic Sulphur-Oxidising Bacteria with Bioleaching Potential: Implications for Pollutant Mobilisation from Mine Waste. Microorganisms 11:222. 10.3390/microorganisms11010222

Pakostova E, McAlary M, Marshall S, et al (2022) Microbiology of a multi-layer biosolid/desulfurized tailings cover on a mill tailings impoundment. J Environ Manage 302:114030. 10.1016/j.jenvman.2021.114030

Peeters C, De Canck E, Cnockaert M, et al (2019) Comparative Genomics of Pandoraea, a Genus Enriched in Xenobiotic Biodegradation and Metabolism. Front Microbiol 10:1–21. 10.3389/fmicb.2019.02556

Pronk JT, Meulenberg R, Hazeu W, et al (1990) Oxidation of reduced inorganic sulphur compounds by acidophilic thiobacilli. FEMS Microbiol Lett 75:293–306. 10.1111/j.1574-6968.1990.tb04103.x

Pyne P, Alam M, Rameez MJ, et al (2018) Homologs from sulfur oxidation (Sox) and methanol dehydrogenation (Xox) enzyme systems collaborate to give rise to a novel pathway of chemolithotrophic tetrathionate oxidation. Mol Microbiol 109:169–191. 10.1111/mmi.13972

Rameez MJ, Pyne P, Mandal S, et al (2020) Two pathways for thiosulfate oxidation in the alphaproteobacterial chemolithotroph Paracoccus thiocyanatus SST. Microbiol Res 320:. 10.1016/j.micres.2019.126345

Rangasamy A, Veeranan J, Pandiyan IG, et al (2014) Early plant growth promotion of maize by various sulfur oxidizing bacteria that uses different thiosulfate oxidation pathway. African J Microbiol Res 8:19–27. 10.5897/ajmr2013.5661

Ray WK, Zeng G, Potters MB, et al (2000) Characterization of a 12-kilodalton rhodanese encoded by glpE of Escherichia coli and its interaction with thioredoxin. J Bacteriol 182:2277–2284. 10.1128/JB.182.8.2277-2284.2000

Rethmeier J, Rabenstein A, Langer M, Fischer U (1997) Detection of traces of oxidized and reduced sulfur compounds in small samples by combination of different high-performance liquid chromatography methods. J Chromatogr A 760:295–302. 10.1016/S0021-9673(96)00809-6

Romero R, Viedma P, Cotoras D (2024) Biooxidation of hydrogen sulfide to sulfur by moderate thermophilic acidophilic bacteria. Biodegradation 35:195–208. 10.1007/s10532-023-10049-y

Sahin N, Tani A, Kotan R, et al (2011) Pandoraea oxalativorans sp. nov., Pandoraea faecigallinarum sp. nov. and Pandoraea vervacti sp. nov., isolated from oxalate-enriched culture. Int J Syst Evol Microbiol 61:2247–2253. 10.1099/ijs.0.026138-0

Satoh H, Odagiri M, Ito T, Okabe S (2009) Microbial community structures and in situ sulfate-reducing and sulfur-oxidizing activities in biofilms developed on mortar specimens in a corroded sewer system. Water Res 43:4729–4739. 10.1016/j.watres.2009.07.035

Schwarz A, Suárez JI, Aybar M, et al (2020) A membrane-biofilm system for sulfate conversion to elemental sulfur in mining-influenced waters. Sci Total Environ 740:. 10.1016/j.scitotenv.2020.140088

Sievert SM, Heidorn T, Kuever J (2000) Halothiobacillus kellyi sp. nov., a mesophilic, obligately chemolithoautotrophic, sulfur-oxidizing bacterium isolated from a shallow-water hydrothermal vent in the Aegean Sea, and emended description of the genus Halothiobacillus. Int J Syst Evol Microbiol 50:1229–1237. 10.1099/00207713-50-3-1229

Spallarossa A, Donahue JL, Larson TJ, et al (2001) Escherichia coli GlpE is a prototype sulfurtransferase for the single-domain rhodanese homology superfamily. Structure 9:1117– 1125. 10.1016/S0969-2126(01)00666-9

Stumm W, Morgan JJ (1995) Aquatic Chemistry: Chemical Equilibria and Rates in Natural Waters, 3rd ed. New York : Wiley, C1996.

Sun X, Kong T, Li F, et al (2022) Desulfurivibrio spp. mediate sulfur-oxidation coupled to Sb(V) reduction, a novel biogeochemical process. ISME J 16:1547–1556. 10.1038/s41396-022-01201-2

Tanabe TS, Dahl C (2022) <scp>HMS-S-S</scp> : A tool for the identification of Sulphur metabolism-related genes and analysis of operon structures in genome and metagenome assemblies. Mol Ecol Resour 22:2758–2774. 10.1111/1755-0998.13642

Tano T, Kitaguchi H, Harada M, et al (1996) Purification and Some Properties of a Tetrathionate Decomposing Enzyme from Thiobacillus thiooxidans. Biosci Biotechnol Biochem 60:224–227. 10.1271/bbb.60.224

Twible LE, Whaley-Martin K, Chen L-X, et al (2024) pH and thiosulfate dependent microbial sulfur oxidation strategies across diverse environments. Front Microbiol 15:. 10.3389/fmicb.2024.1426584

van Vliet DM, von Meijenfeldt FAB, Dutilh BE, et al (2021) The bacterial sulfur cycle in expanding dysoxic and euxinic marine waters. Environ Microbiol 23:2834–2857. 10.1111/1462-2920.15265

Varga D, Horváth AK (2007) Kinetics and Mechanism of the Decomposition of Tetrathionate Ion in Alkaline Medium. Inorg Chem 46:7654–7661. 10.1021/ic700992u

Vigneron A, Cruaud P, Culley AI, et al (2021) Genomic evidence for sulfur intermediates as new biogeochemical hubs in a model aquatic microbial ecosystem. Microbiome 9:1–14. 10.1186/s40168-021-00999-x

Wang J, Cheng Z, Wang J, et al (2023) Temperature and pH on Microbial Desulfurization of Sulfide Wastewater: From Removal Performance to Gene Regulation Mechanism. SSRN Electron J 53:103720. 10.2139/ssrn.4326619

Wang R, Lin JQ, Liu XM, et al (2019) Sulfur oxidation in the acidophilic autotrophic Acidithiobacillus spp. Front Microbiol 10:1–20. 10.3389/fmicb.2018.03290

Wang S, Jiang L, Hu Q, et al (2021) Characterization of Sulfurimonas hydrogeniphila sp. nov., a Novel Bacterium Predominant in Deep-Sea Hydrothermal Vents and Comparative Genomic Analyses of the Genus Sulfurimonas. Front Microbiol 12:1–20. 10.3389/fmicb.2021.626705

Warren LA, Norlund KLI, Bernier L (2008) Microbial thiosulphate reaction arrays: the interactive roles of Fe(III), O 2 and microbial strain on disproportionation and oxidation pathways. Geobiology 6:461–470. 10.1111/j.1472-4669.2008.00173.x

Wasmund K, Mußmann M, Loy A (2017) The life sulfuric: microbial ecology of sulfur cycling in marine sediments. Environ Microbiol Rep 9:323–344. 10.1111/1758-2229.12538

Watanabe T, Kojima H, Umezawa K, et al (2019) Genomes of neutrophilic sulfur-oxidizing chemolithoautotrophs representing 9 proteobacterial species from 8 genera. Front Microbiol 10:1–13. 10.3389/fmicb.2019.00316

Watling HR, Collinson DM, Fjastad S, et al (2014) Column bioleaching of a polymetallic ore: Effects of pH and temperature on metal extraction and microbial community structure. Miner Eng 58:90–99. 10.1016/j.mineng.2014.01.022

Wentzien SW, Sand W (2004) Tetrathionate disproportionation by Thiomonas intermedia K12. Eng Life Sci 4:25–30. 10.1002/elsc.200400007

Whaley-Martin K, Chen L, Nelson TC, et al (2023) O2 partitioning of sulfur oxidizing bacteria drives acidity and thiosulfate distributions in mining waters. Nat Commun 14:2006. 10.1038/s41467-023-37426-8

Whaley-Martin K, Jessen GL, Colenbrander-Nelson T, et al (2019) The Potential Role of Halothiobacillus spp. in Sulfur Oxidation and Acid Generation in Circum-Neutral Mine Tailings Reservoirs. Front Microbiol 10:. 10.3389/fmicb.2019.00297

Whaley-Martin K, Marshall S, Colenbrander-Nelson T, et al (2020) A Mass-Balance Tool for Monitoring Potential Dissolved Sulfur Oxidation Risks in Mining Impacted Waters. Mine Water Environ 39:291–307. 10.1007/s10230-020-00671-0

Whaley-Martin KJ, Chen L-X, Nelson TC, et al (2023b) O2 partitioning of sulfur oxidizing bacteria drives acidity and thiosulfate distributions in mining waters. Nat Commun 14:2006. 10.1038/s41467-023-37426-8

Wood AP, Woodall CA, Kelly DP (2005) Halothiobacillus neapolitanus strain OSWA isolated from “The Old Sulphur Well” at Harrogate (Yorkshire, England). Syst Appl Microbiol 28:746–748. 10.1016/j.syapm.2005.05.013

Yong D, Ee R, Lim YL, et al (2016) Complete genome sequence of Pandoraea thiooxydans DSM 25325T, a thiosulfate-oxidizing bacterium. J Biotechnol 217:51–52. 10.1016/j.jbiotec.2015.11.009

Yu Q, Sun W, Gao H (2021) Thiosulfate oxidation in sulfur-reducing Shewanella oneidensis and its unexpected influences on the cytochrome c content. Environ Microbiol 23:7056–7072. 10.1111/1462-2920.15807

Yu Y, Liu X, Wang H, et al (2014) Construction and Characterization of tetH Overexpression and Knockout Strains of Acidithiobacillus ferrooxidans. J Bacteriol 196:2255–2264. 10.1128/JB.01472-13

Zhang J, Liu R, Xi S, et al (2020) A novel bacterial thiosulfate oxidation pathway provides a new clue about the formation of zero-valent sulfur in deep sea. ISME J 14:2261–2274. 10.1038/s41396-020-0684-5

