## Supplemental Tables 1,2,4,5,6,7 for "Beyond SOx: Mining Sulfur Bacteria Produce Less Acid due to Unidentified Enzyme Pathways"

JAY GORDON

[1]CIVIL AND MINERAL ENGINEERING, UNIVERSITY OF TORONTO, TORONTO, ONTARIO, CANADA

**SI Table 1** Review of Studies in Sulfur Enzyme Literature for Mining Wastewaters

| System Studied | 16S rRNA SOB | Metagenomes/ mRNA SOB | Geochemistry Products/ reactants | Geochemistry: Mass Balance | Reference |
|---|---|---|---|---|---|
| Mine Wastewater | 1 | 1 | 1 | 1 | (Sun et al. 2022) |
| – field | 1 | 1 | 1 | 1 | (Twible et al. 2024) |
| | 1 | 1 | 1 | 0 | (Whaley-Martin et al. 2023a) |
| Mine Wastewater | 1 | 0 | 1 | 1 | (Miettinen et al. 2021) |
| – field & lab mixed communities | 1 | 1 | 1 | 0 | (Whaley-Martin et al. 2019) |
| | 1 | 1 | 1 | 0 | (Camacho et al. 2020c) |
| Mine Wastewater | 0 | 0 | 1 | 0 | (Sun et al. 2022) |
| – lab mixed communities | 0 | 1 | 1 | 1 | (Opara et al. 2023) |
| Mine Wastewater | 1 | 0 | 1 | 0 | (Camacho et al. 2020a) |
| – bioreactors/bioleaching | 1 | Targeted genes | 1 | 0 | (Watling et al. 2014) |
| | 1 | 0 | 1 | 1 | (Li et al. 2020) |
| | 1 | 0 | 1 | 0 | (Schwarz et al. 2020) |
| | 0 | TetH | 1 | 0 | (Romero et al. 2024) |
| | 1 | 0 | 1 | 0 | (Bugaytsova and Lindström 2004) |
| Industrial Wastewater – lab grown mixed communities | 1 | 1 | 1 | 0 | (Chen et al. 2004) |
| Mine Wastewater – SRB | 0 | 0 | 1 | 0 | (Wang et al. 2023) |
| | 1 | 1 | 1 | 0 | (Eckley et al. 2015) |

1 = Data present

0 = Data absent

**SI Table 1 Cont.** Review of Studies in Sulfur Enzyme Literature for Mining Wastewaters

| System Studied | 16S rRNA SOB | Metagenomes/ mRNA SOB | Geochemistry Products/ reactants | Geochemistry: Mass Balance | Reference |
|---|---|---|---|---|---|
| Laboratory SOB Isolates | 1 | 1 | 0 | 0 | (Sun et al. 2022) |
| | 1 | 0 | 1 | 0 | (Chua et al. 2019) |
| | 0 | 1 | 1 | 0 | (Anandham et al. 2010) |
| | 1 | 0 | 1 | 0 | (Peeters et al. 2019) |
| | 1 | SoxB only | 1 | 1 | (Sahin et al. 2011) |
| | 1 | 0 | 1 | 0 | (Anandham et al. 2008) |
| | 1 | 0 | 1 | 0 | (Wood et al. 2005) |
| | 1 | 0 | 1 | 0 | (Sievert et al. 2000) |
| | 1 | 1 | 0 | 0 | (Chen et al. 2004) |
| | 0 | 0 | 1 | 1 | (Arsène-Ploetze et al. 2010) |
| | 0 | functional genes | 0 | 0 | (Wentzien and Sand 2004) |
| Lit Reviews | 1 | 1 | 0 | 0 | (Kappler et al. 2001) |
| | 1 | 1 | 0 | 0 | (Anantharaman et al. 2018) |
| | 1 | no raw data | no raw data | 0 | (van Vliet et al. 2021) |
| | no raw data | no raw data | no raw data | no raw data | (Han and Perner 2015) |
| | 1 | 1 | 0 | 0 | (Hao et al. 2014) |
| | 1 | 1 | 0 | 0 | (Friedrich et al. 2001) |
| | 1 | 1 | 0 | 0 | (Watanabe et al. 2019) |
| | no raw data | no raw data | no raw data | 0 | (Wasmund et al. 2017) |
| | 1 | 1 | 1 | 0 | (Ghosh and Dam 2009) |

1 = Data present

0 = Data absent

**SI Table 2** Potential Enzyme Facilitated SOx and $S_4I$ Pathway Reactions from Current Literature*

| **Part A: Established S Enzyme Pathways** | | | | |
|---|---|---|---|---|
| Pathway | S Gene | Sulfur Substrate Half Reaction | Oxygen as TEA Half Reaction | $\Delta H^+/S$ |
| cSOx | *soxAX, soxYZ* | $S_2O_3^{2-} + \text{SoxYZ-S}^- \rightarrow \text{SoxYZ-S-S-SO}_3^{2-} + 2\ e^-$ | $2\ e^- + 2H^+ + \frac{1}{2}\ O_2 \rightarrow H_2O$ | -1 |
| | *soxB* | $\text{SoxYZ-S}_3O_3^{2-} + H_2O \rightarrow SO_4^{2-} + 2\ H^+ + \text{SoxYZ-S-S}^-$ | NA | 1 |
| | *soxCD* | $\text{SoxYZ-S-S}^- + 3\ H_2O \rightarrow \text{SoxYZ-S-SO}_3^{2-} + 6\ H^+ + 6\ e^-$ | $6\ e^- + 6\ H^+ + 3/2\ O_2 \rightarrow 3\ H_2O$ | 0 |
| | *soxB* | $\text{SoxYZ-S-SO}_3^{2-} + H_2O \rightarrow SO_4^{2-} + 2\ H^+ + \text{SoxYZ-S}^-$ | NA | 1 |
| **cSOx Overall** | ***soxAXBCDYZ*** | $\mathbf{S_2O_3^{2-} + H_2O + 2\ O_2 \rightarrow 2\ SO_4^{2-} + 2\ H^+}$ | | **1** |
| iSOx | *soxAX, soxYZ* | $S_2O_3^{2-} + \text{SoxYZ-S}^- \rightarrow \text{SoxYZ-S-S-SO}_3^{2-} + 2\ e^-$ | $2\ e^- + 2H^+ + \frac{1}{2}\ O_2 \rightarrow H_2O$ | -1 |
| | *soxB* | $\text{SoxYZ-S}_3O_3^{2-} + H_2O \rightarrow SO_4^{2-} + 2\ H^+ + \text{SoxYZ-S-S}^-$ | NA | 1 |
| **iSOx Overall** | ***soxAXBYZ*** | $\mathbf{S_2O_3^{2-} + H_2O + 2\ O_2 \rightarrow 2\ SO_4^{2-} + 2\ H^+}$ | | **0** |
| $S_4I$ – Part 1 | *doxDA*[1] / *tsdAB*[2] | $2\ S_2O_3^{2-} \rightarrow S_4O_6^{2-} + 2\ e^-$ | $2\ e^- + 2\ H^+ + \frac{1}{2}\ O_2 \rightarrow H_2O$ | -0.5 |
| **P1 Overall** | | $\mathbf{2\ S_2O_3^{2-} + 2\ H^+ + \frac{1}{2}\ O_2 \rightarrow S_4O_6^{2-} + H_2O}$ | | **-0.5** |
| $S_4I$ – Part 2 / Other SOI cycling | *tetH*[3] | $S_4O_6^{2-} + H_2O \rightarrow S_2O_3^{2-} + S^0 + SO_4^{2-} + 2\ H^+$ (via DSMSA) | NA | 0.5 |
| | *sdoAB*[4] | $S^0 + 3\ H_2O \rightarrow SO_3^{2-} + 6\ H^+ + 4\ e^-$ | $4\ e^- + 4\ H^+ + O_2 \rightarrow 2\ H_2O$ | 2 |
| | *soeABC / sorAB /abiotic/SUOX* | $SO_3^{2-} + H_2O \rightarrow SO_4^{2-} + 2\ H^+ + 2\ e^-$ | $2\ e^- + 2\ H^+ + \frac{1}{2}\ O_2 \rightarrow H_2O$ | 0 |
| | *SOR*[5] | $4\ \text{R-S-S}/S^0 + 6\ H_2O \rightarrow HS^- + S_2O_3^{2-} + SO_3^{2-} + 11\ H^+ + 6\ e^-$ | $6\ e^- + 6\ H^+ + 3/2\ O_2 \rightarrow 3\ H_2O$ | 5/4 |
| S reduction | *otr/ttrABC*[6] | $S_4O_6^{2-} + 2\ e^- \rightarrow 2\ S_2O_3^{2-}$ | NA, S is e- acceptor | |

NA = e- acceptor not required for this reaction (example: decomposition, double replacement, disproportionation)

**SI Table 2 cont.** Potential Enzyme Facilitated SOx and $S_4I$ Pathway Reactions from Current Literature*

| Part B: Speculative S Enzyme Pathways | | | |
|---|---|---|---|
| Pathway | S Gene | Sulfur Substrate Half Reaction | Oxygen as TEA Half Reaction |
| $S_4I$ – Part 2 / Other SOI cycling | *pdo/TST*[7] | $S_2O_3^{2-} + 3 H_20 \rightarrow 2 SO_3^{2-} + 6 H^+ + 4 e^-$ | $4 e^- + 4 H^+ + O_2 \rightarrow 2 H_2O$ |
| | *tetH*[8];<br>*spontaneous* | $S_4O_6^{2-} + H_2O \rightarrow {}^-S\text{-}S\text{-}SO_3^- + SO_4^{2-} + 2 H^+$;<br>${}^-S\text{-}S\text{-}SO_3^- \rightarrow S_2O_3^{2-} + S^0$ **or**<br>$2\ {}^-S\text{-}S\text{-}SO_3^- \rightarrow {}^-S\text{-}S\text{-}S\text{-}S\text{-}SO_3^- + SO_3^{2-}$<br>$2\ {}^-S\text{-}S\text{-}S\text{-}S\text{-}SO_3^- \rightarrow S_9O_3^{2-} + SO_3^{2-}$<br>$S_9O_3^{2-} \rightarrow SO_3^{2-} + S_8$ | NA<br>NA<br>NA<br>NA<br>NA |
| | *tetH*[9] | $2 S_4O_6^{2-} + H_20 \rightarrow S_2O_3^{2-} + S_5O_6^{2-} + SO_4^{2-} + 2 H^+$ | NA |
| | *tetH*[10] | $4 S_4O_6^{2-} + 5 H_20 \rightarrow 7 S_2O_3^{2-} + 2 SO_4^{2-} + 10 H^+$ | NA |
| | *ThdT*[11] | $2 S_2O_3^{2-} \rightarrow S_4O_6^{2-} + 2 e^-$ /<br>$GSH + S_4O_6^{2-} \rightarrow GSSSSO_3^- + SO_3^{2-} + H^+$ | $2 e^- + 2 H^+ + \frac{1}{2} O_2 \rightarrow H_2O$<br>NA |
| | *soxB*<br>*soxCD*<br>*soxB*[12] | $GSSSSO_3^- + H_20 \rightarrow GSSS^- + SO_4^{2-} + 2 H^+$<br>$GSSS^- + 3 H_20 \rightarrow GSSSO_3^- + 6 H^+ + 6 e^-$<br>$GSSSSO_3^- + H_20 \rightarrow GSSS^- + SO_4^{2-} + 2 H^+$ | NA<br>$6 e^- + 6 H^+ + 3/2 O_2 \rightarrow 3 H_2O$<br>NA |
| | *soxB*[13] | $S_4O_6^{2-} + 2 H_2O \rightarrow R\text{-}SS/S^0 + 2 SO_4^{2-} + 4 H^+ + 2 e^-$ | $2 e^- + 2 H^+ + \frac{1}{2} O_2 \rightarrow H_2O$ |
| | *trithionate hydrolase*[14] | $S_3O_6^{2-} + H_2O \rightarrow S_2O_3^{2-} + SO_4^{2-} + 2 H^+$ | NA |
| | *SOR*[15] | $7 R\text{-}S\text{-}S/S^0 + 3 H_2O + 8 e^- \rightarrow 6 HS^- + SO_3^{2-}$ | NA |
| | *gplE*[16] | $S_2O_3^{2-} + CN^- \rightarrow SO_3^{2-} + SCN^-$ | NA |
| | *tetH*[17] | $S_5O_6^{2-} + H_2O \rightarrow 2S^0 + S_2O_3^{2-} + SO_4^{2-} + 2 H^+$ | NA |

**SI Table 2 cont.** Potential Enzyme Facilitated SOx and $S_4I$ Pathway Reactions from Current Literature*

| **Part B: Speculative S Enzyme Pathways** | | | |
|---|---|---|---|
| Pathway | S Gene | Sulfur Substrate Half Reaction | Oxygen as TEA Half Reaction |
| | *Not matched to gene* [18] | $S_3O_3^{2-} + S_4O_6^{2-} \rightarrow S_2O_3^{2-} + S_5O_6^{2-}$<br>$2\ S_3O_3^{2-} + S_4O_6^{2-} \rightarrow 2\ S_2O_3^{2-} + S_6O_6^{2-}$<br>$S_3O_3^{2-} + S_5O_6^{2-} \rightarrow S_2O_3^{2-} + S_6O_6^{2-}$<br>$4\ S_3O_3^{2-} \rightarrow S_8 + 4\ SO_3^{2-}$ | NA |
| $S_4I$ – Part 2 / Other SOI cycling | *Abiotic* [19] | $S_4O_6^{2-} + SO_3^{2-} \rightarrow S_3O_6^{2-} + S_2O_3^{2-}$ | NA |
| | *Abiotic* [20] | $1/8\ S_8 + SO_3^{2-} \rightarrow S_2O_3^{2-}$ | NA |
| | *Not matched to gene* [21] | $4\ S_4O_6^{2-} + 4\ H_2O \rightarrow 6\ S_2O_3^{2-} + S_3O_6^{2-} + SO_4^{2-} + 8\ H^+$ | NA, disproportionation |
| S reduction | *ttrABC* [22] | $S_3O_6^{2-} + 2\ e^- \rightarrow S_2O_3^{2-} + SO_3^{2-}$ | NA, S is e- acceptor |
| | *tsrABC* [23] *(aka PhsABC)* | $S_2O_3^{2-} + H^+ + 2\ e^- \rightarrow HS^- + SO_3^{2-}$ | NA, S is e- acceptor |

**S compounds:** $S_8$ = elemental sulfur / cyclooctasulfur; R-SS$^-$ = cysteine-bound persulphide; $^-$S-S$_n$-S$^-$ / $S_n^{2-}$ = inorganic polysulphide; $S_3O_3^{2-}$ /$^-$S-S-SO$_3^-$ = disulfane monosulfonic acid; GS–S–S–SO$_3^-$ = glutathione:sulfodisulfane adduct**;** DSMSA = Disulfane monosulfonic acid; PT = pentathionate; TSMSA = tetrasulfane monosulfonic acid; OSMSA = octosulfane monosulfonic acid **S metabolism enzymes:** Dox, a type of thiosulfate:quinone oxidoreductase (TQO); GplE, thiosulfate:cyanide sulfurtranserase/thiosulfate sulfurtransferase; Otr – octoheme tetrathionate reductase; Pdo, persulphide dioxygenase; Sdo, sulfur dioxygenase; Soe, sulfite- oxidizing enzyme; SOR, sulfur oxygenase/reductase; Sor, sulfite:acceptor oxidoreductase; Sox, sulfur-oxidizing multienzyme complex; TdhT - thiol dehydrotransferase; TetH / TTH / 4THase, tetrathionate hydrolase; Tsd, thiosulfate dehydrogenase; Tsr/Phs, thiosulfate reductase; Ttr, tetrathionate reductase; TST, thiosulfate:cyanide sulfur transferase.

**SI Table 2 cont.** Potential Enzyme Facilitated SOx and $S_4I$ Pathway Reactions from Current Literature*

**References**

| Footnotes | Study |
|---|---|
| 1 | (Müller et al. 2004) |
| 2 | (Denkmann et al. 2012; Brito et al. 2015; Jenner et al. 2019; Li et al. 2020; Rameez et al. 2020; Yu et al. 2021; Du et al. 2022) |
| 3 | (Kanao et al. 2007), (Meulenberg et al. 1992b) (Tanabe and Dahl 2022) (De Jong et al. 1997) |
| 4 | (Wang et al. 2019; Zhang et al. 2020) |
| 5 | (Kletzin 1992; Ghosh and Dam 2009) |
| 6 | (Hensel et al. 1999) |
| 7 | (Vigneron et al. 2021) |
| 8 | (Dahl 2005; Gwak et al. 2022) |
| 9 | (Bugaytsova and Lindström 2004) |
| 10 | (Tano et al. 1996) |
| 11 | (Pyne et al. 2018) |
| 12 | (Pyne et al. 2018) |
| 13 | (Cai et al. 2022) |
| 14 | (Meulenberg et al. 1992a), (Anandham et al. 2008),(Lu and Kelly 1988) (Pronk et al. 1990) |
| 15 | (Tanabe and Dahl 2022) |
| 16 | (Ray et al. 2000; Spallarossa et al. 2001) |
| 17 | (De Jong et al. 1997) |
| 18 | (Wentzien and Sand 2004) |
| 19 | (Hinsley and Berks 2002) |
| 20 | (Hinsley and Berks 2002) |
| 21 | (Wentzien and Sand 2004) |
| 22 | (Hinsley and Berks 2002) |
| 23 | (Heinzinger et al. 1995; Hinsley and Berks 2002; Haja et al. 2020) |

**SI Table 3** Complete Geochemical Data from 16 Microcosms

- See separate file: "SI_Table_3_Complete_Geochemical_Data_JG.csv"

**SI Table 4** Relative Abundance of Bacteria Genera in the 16 Mesocosms

| **Microcosm** | **Treatment** | **Time** | **Exp** | ***Acinetobacter*** | ***Allorhizobium-Neorhizobium-Pararhizobium-Rhizobium*** | ***Brucella*** | ***Delftia*** | ***Halothiobacillus*** | ***Massilia*** | ***Pandoraea*** | ***Pseudomonas*** | ***Sphingobium*** | ***Stenotropho-monas*** | ***Thiomonas*** | ***Xanthobacter*** |
|---|---|---|---|---|---|---|---|---|---|---|---|---|---|---|---|
| All | 2 m SOB + Tetra | $t_0$ | a | 0.13 | 0.00 | 0.00 | 85.64 | 0.52 | 0.00 | 11.69 | 1.16 | 0.01 | 0.03 | 0.75 | 0.00 |
| M3 | 2 m SOB (No SOI) R1 | $t_{end}$ | a | 0.01 | 0.00 | 0.00 | 90.13 | 0.54 | 0.01 | 8.76 | 0.17 | 0.00 | 0.01 | 0.32 | 0.00 |
| M4 | 2 m SOB (No SOI) R 2 | $t_{end}$ | a | 0.00 | 0.00 | 0.00 | 0.07 | 78.08 | 0.00 | 20.73 | 0.37 | 0.08 | 0.02 | 0.32 | 0.32 |
| M5 | 2 m SOB + Tetra R1 | $t_{end}$ | a | 0.01 | 0.00 | 0.00 | 0.03 | 90.11 | 0.00 | 8.79 | 0.16 | 0.05 | 0.02 | 0.52 | 0.31 |
| M6 | 2 m SOB + Tetra R2 | $t_{end}$ | a | 0.00 | 0.00 | 0.00 | 0.00 | 99.30 | 0.00 | 0.11 | 0.00 | 0.00 | 0.00 | 0.49 | 0.09 |
| All | 10 m SOB + Tetra | $t_0$ | b | 0.00 | 0.05 | 0.00 | 0.13 | 0.41 | 0.00 | 73.99 | 19.74 | 4.97 | 0.05 | 0.14 | 0.46 |
| M7 | 10 m SOB (No SOI) R1 | $t_{end}$ | b | 0.00 | 0.03 | 0.00 | 29.09 | 0.55 | 0.00 | 12.26 | 55.09 | 1.84 | 0.78 | 0.13 | 0.06 |
| M8 | 10 m SOB (No SOI) R2 | $t_{end}$ | b | 0.04 | 0.01 | 0.00 | 2.54 | 89.07 | 0.00 | 3.44 | 0.29 | 0.04 | 0.06 | 4.17 | 0.00 |
| M9 | 10 m SOB + Tetra R1 | $t_{end}$ | b | 0.03 | 0.00 | 0.00 | 0.42 | 91.78 | 0.00 | 3.28 | 0.25 | 0.04 | 0.24 | 3.77 | 0.00 |
| M10 | 10 m SOB + Tetra R2 | $t_{end}$ | b | 0.00 | 0.00 | 0.00 | 0.00 | 89.69 | 0.00 | 1.65 | 0.06 | 0.00 | 0.00 | 8.59 | 0.00 |
| M13/ M14 | 2 m SOB + Thio | $t_0$ | c | 0.04 | 0.00 | 0.00 | 0.00 | 96.67 | 0.00 | 0.15 | 0.02 | 0.00 | 0.00 | 2.14 | 0.97 |
| M11/ M12 | 10 m SOB + Thio | $t_0$ | c | 0.16 | 0.00 | 0.00 | 0.00 | 94.17 | 0.00 | 0.15 | 0.05 | 0.00 | 0.01 | 3.06 | 1.48 |
| M11 | 2 m SOB + Thio R1 | $t_{end}$ | c | 0.04 | 2.08 | 0.01 | 0.00 | 59.99 | 0.00 | 32.64 | 3.86 | 0.22 | 0.22 | 0.37 | 0.54 |
| M12 | 2 m SOB + Thio R2 | $t_{end}$ | c | 0.00 | 0.00 | 0.77 | 0.00 | 94.97 | 0.00 | 2.97 | 0.01 | 0.00 | 0.23 | 0.70 | 0.32 |
| M13 | 10 m SOB + Thio R1 | $t_{end}$ | c | 0.00 | 0.43 | 2.14 | 0.00 | 2.99 | 0.00 | 3.53 | 88.18 | 0.07 | 2.37 | 0.03 | 0.20 |
| M14 | 10 m SOB + Thio R2 | $t_{end}$ | c | 0.00 | 0.00 | 0.00 | 0.00 | 87.93 | 0.00 | 11.50 | 0.01 | 0.00 | 0.04 | 0.12 | 0.34 |

- For workflow from raw 16S gene sequencing data with OTUs listed, see "Chapter 4 Supplementary 16S rRNA data (curation steps)" at the following link: http://128.100.14.155:8088/share.cgi?ssid=204a2ab4254d4911874cd20953dfb1ed.

**SI Table 5** Frequency of metagenomes of SOB containing one or more copy/ies of each S gene

| | Potential $S_4I$ | | | | | | | SOx | | | | | | rDSR | | | | | | | | | | | S Uptake |
|---|---|---|---|---|---|---|---|---|---|---|---|---|---|---|---|---|---|---|---|---|---|---|---|---|---|
| | ***tsdA*** | ***doxDA*** | ***sdo*** | ***TST*** | ***sor*** | ***SUOX*** | ***ttrABC*** | ***soxA*** | ***soxB*** | ***soxC*** | ***soxX*** | ***soxY*** | ***soxZ*** | ***dsrC*** | ***dsrA*** | ***dsrB*** | ***dsrE*** | ***dsrF*** | ***dsrH*** | ***dsrC*** | ***dsrMKJOP*** | ***aprAB*** | ***sat*** | ***qmoABC*** | ***sqr*** |
| *Halothiobacillus*[1] (Ox Res 10 m) | 1 | 0 | 1 | 1 | 1 | NA | 0 | 1 | 1 | 1 | 1 | 1 | 1 | 0 | 0 | 0 | 0 | 0 | 0 | 0 | 0 | 0 | 0 | 0 | 1 |
| *Halothiobacillus* (Ox Res 2 m) | 1 | 0 | 1 | 1 | 0 | NA | 0 | 1 | 1 | 1 | 1 | 1 | 1 | 0 | 0 | 0 | 0 | 0 | 0 | 0 | 0 | 0 | 0 | 0 | 1 |
| *Thiomonas* (Ox Res Average) | 1 | 0 | 1 | 1 | 0 | NA | 0 | 1 | 1 | 1 | 1 | 1 | 1 | 0 | 0 | 0 | 0 | 0 | 0 | 0 | 0 | 0 | 0 | 0 | 1 |
| *Pandoraea* (Lit review[2]) | - | - | - | 1 | - | 1 | - | 1 | 1 | - | 1 | 1 | 1 | - | - | - | - | - | - | - | - | - | - | - | 1 |
| *Pandoraea* (Flin Flon*) | 0 | 0 | 1 | 1 | 0 | NA | 0 | 1 | 1 | 1 | 1 | 1 | 1 | 0 | 0 | 0 | 0 | 0 | 0 | 0 | 0 | 0 | 0 | 0 | 1 |

NA – data not available from analysis

[1]The following genes were not detected in *Halothiobacillus:* cysA, cysC, cysD, cysl, cysP, cysW, cycU, psrA, psrB, phsA, sudA, sudB, hdrB1, hdrA1, hdrB2, hdrC2, asrA_asrB, asrC, sir, sreC and fsr; hdrA2 detected in *Thiomonas* but not *Halothiobacillus*; cysJ detected in *Halothiobacillus* but not *Thiomonas.*

[2] *Pandoraea* lit review based on a comparison of over 68 metagenomes (Wang et al. 2019). *Pandoraea* metagenomes also contain: cysA, cysD, cysE, cysJ , cysN, cysP, cysU dmdC, dmdD, ssuA, ssuB, ssuC, ssuD, ssuE, metX, metZ, tauB, tauC (Anandham et al. 2008, 2010; Rangasamy et al. 2014; Yong et al. 2016; Peeters et al. 2019).

Note: *Halothiobacillus* spp. metagenome assembled genomes for the initial communities used in these experiments (2017 2m and 10 m) also contain the ability to reduce nitrite to ammonia (*nirB*), but lack the capacity to reduce nitrate to nitrite (*narGHIJ* / *napAB*) (Peeters et al. 2019). *Thiomonas* spp. metagenomes have the capacity to reduce nitrate (*narGHIJ*) but not nitrite (*nirBSK*) (Whaley-Martin et al. 2023b). Of the 68 *Pandoraea* genomes in the literature review, > 96% have the ability to reduce nitrite to ammonia (*nirB*), but only 25 % contain one of the genes required to reduce nitrate to nitrite ((*narG*), (Whaley-Martin et al. 2023b)).

**SI Table 6** Calculation of Approximate Oxygen Utilization Rate (OUC) in Anoxic Microcosms

**Depth of Oxygen Penetration in SOB biofilms with *Halothiobacillus***

Microbial community structures and in situ sulfate-reducing and sulfur-oxidizing activities in biofilms developed on mortar specimens in a corroded sewer system

10.1016/j.watres.2009.07.035

| | | |
|---|---|---|
| Depth of Oxygen penetration (approx.) | 1000 | um |
| | 1 | mm |

**Approximate OUR**

Growth kinetics of hydrogen sulfide oxidizing bacteria in corroded concrete from sewers

10.1016/j.jhazmat.2011.03.005

**Range of oxygen utilization rates in an unstirred batch reactor (250 mg flask)**

| | | |
|---|---|---|
| OUR min | 0 | g m^-3 hr^-1 |
| OUR max | 14 | g m^-3 hr^-1 |
| OUR mean (estimated) | 10 | g m^-3 hr^-1 |
| **Surface area of media** | pi*r^2 - flask @ 6 L depth - pi*r^2 DO probe | |
| At t=0 (6 L depth) | 98 | cm^2 |
| At t=end (2 L depth) | 374 | cm^2 |
| **Volume of oxygen exposed media** | 374 | mm*cm*cm |
| | 3.74E-05 | m^3 |
| OUR * area mean (g m^-3 hr^-1*m^3) | 3.74E-04 | g/hr |
| OUR * area max | 5.24E-04 | g/hr |
| Molar Mass of O2 | 32 | g/mol |
| **Moles of Oxygen mean** | 1.17E-05 | mol/hr |
| | 1.17E+01 | umol/hr |
| **Mol of Oxygen max** | 1.64E-05 | mol/hr |
| | 1.64E+01 | umol/hr |
| **Volume of media in flask (mean)** | 4 | L |
| Mean OUR | 2.9 | uM/hr |
| Max OUR | 4.1 | uM/hr |
| **Duration of Suboxic Portion of Exp c** | 162 | hr |
| **Max Oxygen Consumption** | 0.66 | mM |

**SI Table 7** Calculation of Approx. Thiosulfate Diffusion Distance in Suboxic Microcosms

| | | | |
|---|---|---|---|
| **Volume of media in flask** | | | |
| Vmax | | 2000 | cm^3 |
| Vmean | | 4000 | cm^3 |
| Vmin | | 6000 | cm^3 |
| **Flask as a Cone** | **The Volume of Frustum of Cone=1/3 πH (R2 +Rr+r2 )** | | |
| radius (bottom, R) | | 12 | cm |
| radius (surface of water, r)max | | 6 | cm |
| radius (surface of water, r)mean | | 8 | cm |
| radius (surface of water, r)min | | 10 | cm |
| Solve for H (depth) | V/(1/3*pi*(R^2+Rr+r^2)) | | |
| **Depth of media in flask** | | | |
| dmax | | 23 | cm |
| dmean | | 13 | cm |
| **dmin** | | **5** | **cm** |
| **The Stokes-Einstein Law** | $\mathbf{t=x^2/2D}$ (Peeters et al. 2019) | | |
| **Symbol** | **Variable** | | **Units** |
| t | Time | | s |
| D | diffusion coefficient of a solute in a free solution | | cm^2/s |
| x | mean distance traveled by diffusing solute with time | | cm |
| t = time anoxic | | 162 | hrs |
| Conversion s/hour | | 3600 | s/hr |
| t = time anoxic | | 583200 | s |
| **The Stokes-Einstein Law** | $\mathbf{x^2=2*t*D}$ | | |
| Solve for x, using Dmean | 1.00E-05 | | cm^2/s |
| x | | 3.4 | cm |
| D (for sulfate at 25 degrees) | | 1.07E-05 | cm^2/s |
| x | | 3.5 | cm |
| **D chlorine (fast, small anion)** | | **2.03E-05** | **cm^2/s** |
| **x** | | **4.9** | **cm** |
| D glucose (slow, large molecule) | | 5.00E-06 | cm^2/s |
| x | | 0.19 | cm |

$$\mathbf{x_{max} = d_{min}}$$

Therefore, the highest diffusion distance for thiosulfate ($x_{max}$) was only equal to the depth of the media after the final sampling ($d_{min}$). Prior to final sampling, diffusion would be insufficient to allow contact between the thiosulfate and the dissolved oxygen at the surface of the microcosms.

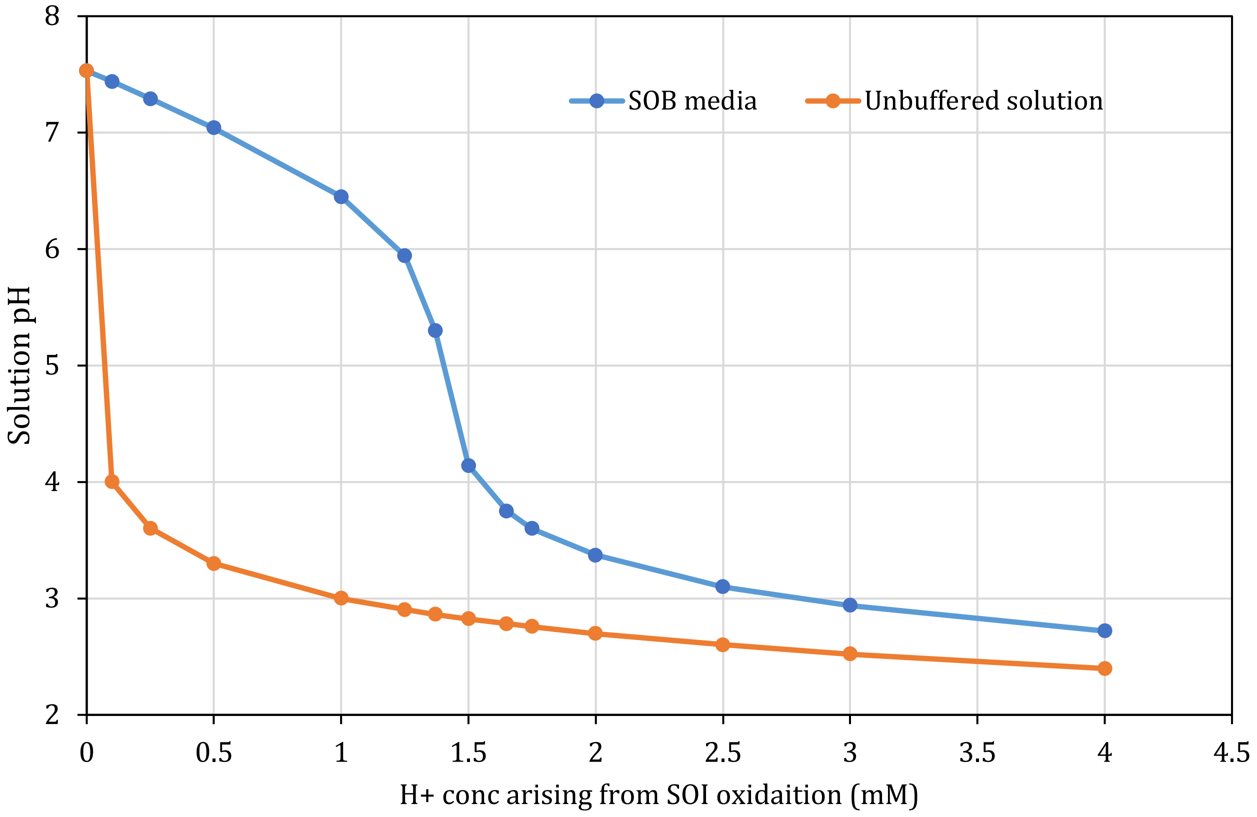

**SI Fig 1** The impacts of carbonate and phosphate buffering in the SOB media (blue line) compared to a solution of unbuffered SOB media (orange line), as calculated via Visual MINTEQ 4.0 (in consultation with Dr. Simon Apte, personal correspondence). The x-axis plots the concentration of protons released into solution via SOI oxidation, and the y-axis indicates the pH which would be observed in buffered or unbuffered media as a result of the respective increases in proton concentration.

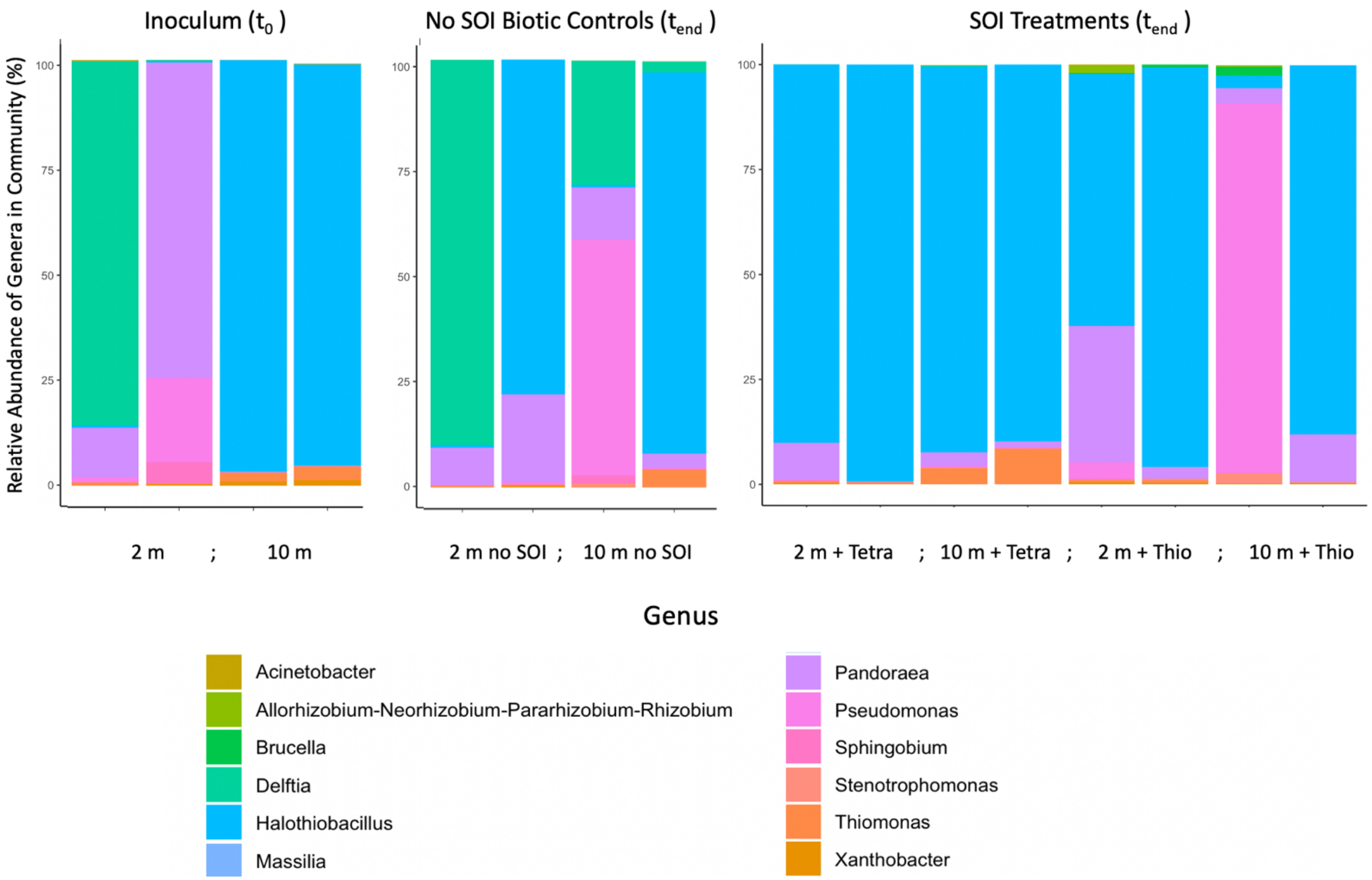

**SI Fig 2** 16S rRNA gene sequencing of microbial communities indicates that the initial microbial communities from 2 m and 10 m ($T_0$) were dominated by the sulfur-oxidizing genera *Halothiobacillus* and *Pandoraea*, and these two genera remained dominant in the SOB and S substrate-amended microcosms (SOI Treatments Tend: $M_5$, $M_6$, $M_9$, $M_{10}$, and $M_{11}$–$M_{14}$), while the heterotrophs *Delftia* and *Pseudomonas* grew in abundance in the microcosm that did not receive S amendments (No SOI Biotic Controls $T_{end}$: $M_3$, $M_4$, $M_7$, and $M_8$).

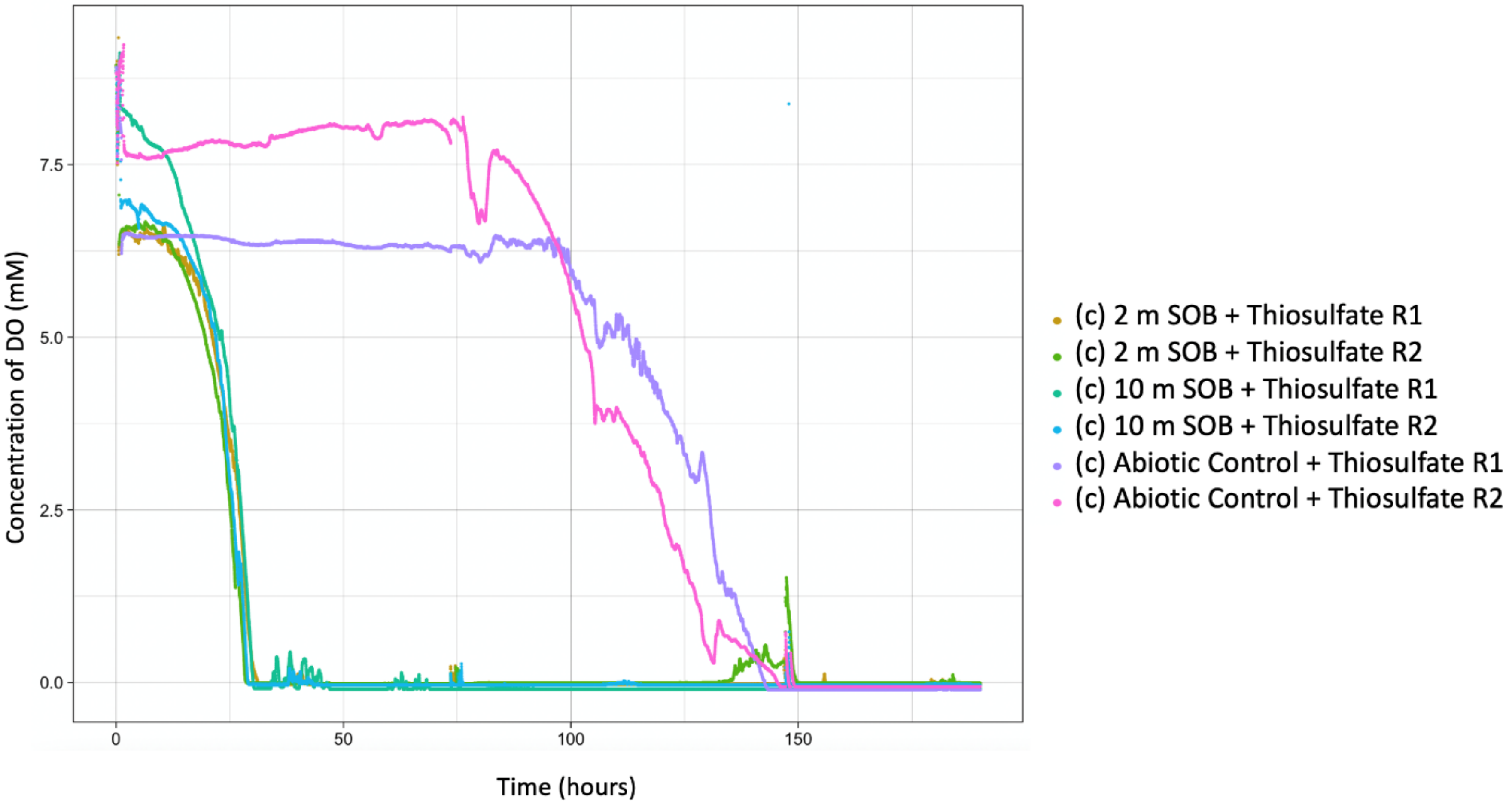

**SI Fig 3** Dissolved oxygen (DO) concentrations in Experiment C demonstrate that the microcosms became suboxic at ~30 hours and maintained suboxic conditions through the remaining 170 hours of the experiment. Probe measurements from 1 cm below the surface of the microcosm also indicated suboxia. (Oxygen loss in the abiotic controls at 100 hours likely indicates contamination of samples with non-sulfur-oxidizing bacteria [SOB] from the atmosphere midway through the experiment.)
